# Antibiotic potentiation and inhibition of cross-resistance in pathogens associated with cystic fibrosis

**DOI:** 10.1101/2023.08.02.551661

**Authors:** Nikol Kadeřábková, R. Christopher D. Furniss, Evgenia Maslova, Kathryn E. Potter, Lara Eisaiankhongi, Patricia Bernal, Alain Filloux, Cristina Landeta, Diego Gonzalez, Ronan R. McCarthy, Despoina A.I. Mavridou

**Affiliations:** Department of Molecular Biosciences, The University of Texas at Austin, Austin, 78712, Texas, USA; John Ring LaMontagne Center for Infectious Diseases, The University of Texas at Austin, Austin, 78712, Texas, USA; Centre for Bacterial Resistance Biology, Department of Life Sciences, Imperial College London, London, SW7 2AZ, UK; Division of Biosciences, Department of Life Sciences, College of Health and Life Sciences, Brunel University London, Uxbridge, UB8 3PH, UK; Departamento de Microbiología, Facultad de Biología, Universidad de Sevilla, Seville, 41012, Spain; Singapore Centre for Environmental Life Sciences Engineering, Nanyang Technological University, 637551, Singapore; School of Biological Sciences, Nanyang Technological University, 639798, Singapore; Lee Kon Chian School of Medicine, Nanyang Technological University, 636921, Singapore; Department of Biology, Indiana University, Bloomington, Indiana, 47405, USA; Laboratoire de Microbiologie, Institut de Biologie, Université de Neuchâtel, Neuchâtel, 2000, Switzerland

**Keywords:** antimicrobial resistance, antibiotic potentiation, cross-resistance, polymicrobial communities, cystic fibrosis, Gram-negative bacterial pathogens, protein homeostasis

## Abstract

Critical Gram-negative pathogens, like *Pseudomonas*, *Stenotrophomonas* and *Burkholderia*, have become resistant to most antibiotics. Complex resistance profiles together with synergistic interactions between these organisms increase the likelihood of treatment failure in distinct infection settings, for example in the lungs of cystic fibrosis (CF) patients. Here, we discover that cell envelope protein homeostasis pathways underpin both antibiotic resistance and cross-protection in CF-associated bacteria. We find that inhibition of oxidative protein folding inactivates multiple species-specific resistance proteins. Using this strategy, we sensitize multidrug-resistant *Pseudomonas aeruginosa* to β-lactam antibiotics and demonstrate promise of new treatment avenues for the recalcitrant emerging pathogen *Stenotrophomonas maltophilia*. The same approach also inhibits cross-protection between resistant *S. maltophilia* and susceptible *P. aeruginosa*, allowing eradication of both commonly co-occurring CF-associated organisms. Our results provide the basis for the development of next-generation strategies that target antibiotic resistance, while also impairing specific interbacterial interactions that enhance the severity of polymicrobial infections.

## INTRODUCTION

Antimicrobial resistance (AMR) is one of the most significant threats to health systems worldwide [1]. Since the end of the “golden age” of antibiotic discovery in the 1970’s, very few new antimicrobial agents have entered the clinic, and most of those that have gained approval are derivatives of existing antibiotic classes [2–4]. Meanwhile, resistance to useful antibiotics is continuously rising, resulting in more than 1.3 million deaths annually [5]. In addition to the undeniable surge of resistance, it is becoming apparent that intra– and inter-species interactions also play a role in AMR and its evolution [6], ultimately posing additional challenges during antibiotic treatment. This necessitates not only the development of novel antimicrobials and strategies that will expand the lifespan of existing antibiotics, but also the implementation of approaches that will address the polymicrobial nature of most infections.

Antibiotic resistance is most commonly evaluated by testing bacterial strains in monoculture. Nonetheless, the majority of clinical infections contain multiple species whose coexistence in complex pathobionts often limits our treatment options. This is of particular importance for recalcitrant infections such as the polymicrobial communities found in the lungs of cystic fibrosis (CF) patients. CF lung infections have become a paradigm for chronic infectious diseases that result in poor quality of life and early patient mortality [7]. Such infections are dominated by highly resistant opportunistic pathogens, including, but not limited to, *Pseudomonas aeruginosa*, *Staphylococcus aureus*, species and strains belonging to the *Burkholderia* complex, and *Stenotrophomonas maltophilia* [8]. Most of these organisms carry an array of resistance mechanisms, like efflux pumps, atypical lipopolysaccharide structures, and β-lactamase enzymes. Their co-occurrence in the CF lung leads to treatment challenges since common clinical care options for one pathogen are not necessarily compatible with the antibiotic susceptibility profiles of other species that are present. For example, on the one hand, *P. aeruginosa* is the most prevalent organism in CF lung infections and its treatment, especially during pulmonary exacerbation episodes, relies heavily on β-lactam compounds [8]. On the other hand, CF microbiomes are increasingly found to encompass *S. maltophilia* [8–10], a globally distributed opportunistic pathogen that causes serious nosocomial respiratory and bloodstream infections [11–13]. *S. maltophilia* is one of the most prevalent emerging pathogens [12] and it is intrinsically resistant to almost all antibiotics, including β-lactams like penicillins, cephalosporins and carbapenems, as well as macrolides, fluoroquinolones, aminoglycosides, chloramphenicol, tetracyclines and colistin. As a result, the standard treatment option for lung infections, i.e., broad-spectrum β-lactam antibiotic therapy, is rarely successful in countering *S. maltophilia* [13,14], creating a definitive need for approaches that will be effective in eliminating both pathogens.

The lack of suitably broad antibiotic regimes able to simultaneously eradicate all pathogens present in specific infection settings is not the only challenge when treating polymicrobial communities. Bacterial interactions between antibiotic-resistant and antibiotic-susceptible bacteria can add to this problem by adversely affecting antibiotic drug sensitivity profiles of organisms that should be treatable [6]. In particular, some antibiotic resistance proteins, like β-lactamases, which decrease the quantities of active drug present, function akin to common goods, since their benefits are not limited to the pathogen that produces them but can be shared with the rest of the bacterial community. This means that their activity enables pathogen cross-resistance when multiple species are present [15,16], something that was demonstrated in recent work investigating the interactions between pathogens that naturally co-exist in CF infections. More specifically, it was shown that in laboratory co-culture conditions, highly drug-resistant *S. maltophilia* strains actively protect susceptible *P. aeruginosa* from β-lactam antibiotics [15]. Moreover, this cross-protection was found to facilitate, at least under specific conditions, the evolution of β-lactam resistance in *P. aeruginosa* [17]. The basis of such interactions could be exploited during the design of novel therapeutic strategies, since targeting appropriate resistance enzymes will not only render their producers susceptible to existing drugs but should also impair their capacity to protect co-existing antibiotic-susceptible strains.

Protein homeostasis in the Gram-negative cell envelope, and in particular the formation of disulfide bonds by the thiol oxidase DsbA [18–22], is essential for the function of many resistance proteins [23]. Oxidative protein folding occurs post-translationally, after translocation of the nascent polypeptide to the periplasm through the general secretion (Sec) system [24]. There, disulfide bond formation assists the assembly of 40% of the cell-envelope proteome [25,26], promotes the biogenesis of virulence factors [27,28], controls the awakening of bacterial persister cells [29], and underpins the function of resistance determinants, including enzymes for which we do not currently have inhibitor compounds, such as metallo-β-lactamases [30]. Here, we reveal the potential of targeting proteostasis pathways, such as disulfide bond formation, as a strategy against pathogens commonly associated with highly resistant polymicrobial infections. Using this approach, we incapacitate species-specific resistance proteins in CF-associated bacteria and simultaneously abrogate protective effects between pathogens that coexist in these infections. Our results demonstrate that such strategies generate compatible treatment options for recalcitrant CF pathogens and, at the same time, eradicate interspecies interactions that impose additional challenges during antibiotic treatment in complex infection settings.

## RESULTS

### Species-specific cysteine-containing β-lactamases depend on oxidative protein folding

#### β-Lactamase activity

To investigate the potential of targeting disulfide bond formation as a strategy to overcome resistance mechanisms in challenging pathogens, we chose to primarily explore β-lactamases that are produced by bacteria intimately associated with CF lung infections. DsbA dependence has been previously shown for a handful of such enzymes [23], like the chromosomally-encoded class B3 metallo-β-lactamase L1-1 from *S. maltophilia* (Table S1), which contributes significantly to AMR in this organism [13], as well as β-lactamases from the GES and OXA families, which are broadly disseminated, but commonly found in *P. aeruginosa* [31,32]. Here, we selected six clinically important β-lactamases from different Ambler classes (classes A, B and D) that are exclusively encoded either by *P. aeruginosa* or by the *Burkholderia* complex. The *P. aeruginosa* enzymes (BEL-1, CARB-2, AIM-1, and OXA-50) are all phylogenetically distinct, while the *Burkholderia* β-lactamases (BPS-1m and BPS-6) belong to the same phylogenetic class (File S1). Class A, C, and D β-lactamases, like the BPS-6 (class A) and OXA-50 (class D) enzymes investigated here, are serine-dependent hydrolases. Serine β-lactamases are structurally related to penicillin binding proteins, which have a major role in the synthesis of the peptidoglycan [33]. By contrast, class B enzymes are evolutionary distinct and rely on one or two Zn^2+^ ions for catalytic activity [30,34]. In addition to belonging to different phylogenetic classes, the selected enzymes have different numbers of cysteines, display varied hydrolytic activities, can be both resident on the chromosome or on mobile genetic elements, and have diverse inhibitor susceptibility profiles (Table S1).

We expressed all six β-lactamases in the *Escherichia coli* K-12 strain MC1000 and its isogenic *dsbA* deletion mutant. This strain background was selected because it has been traditionally used in oxidative protein folding studies [35–38] and it lacks endogenous β-lactamase enzymes or any other mechanisms that could contribute to antibiotic resistance. We recorded β-lactam minimum inhibitory concentration (MIC) values for each enzyme in both strain backgrounds. We found that expression of all test enzymes in the *dsbA* mutant background resulted in markedly reduced MICs for at least one β-lactam antibiotic (Fig. 1 and File S2A), compared to the MICs recorded in the wild-type *E. coli* strain; only differences larger than 2-fold were considered. These results indicate that the presence of DsbA is important for the function of all tested resistance proteins.

**Figure 1.**
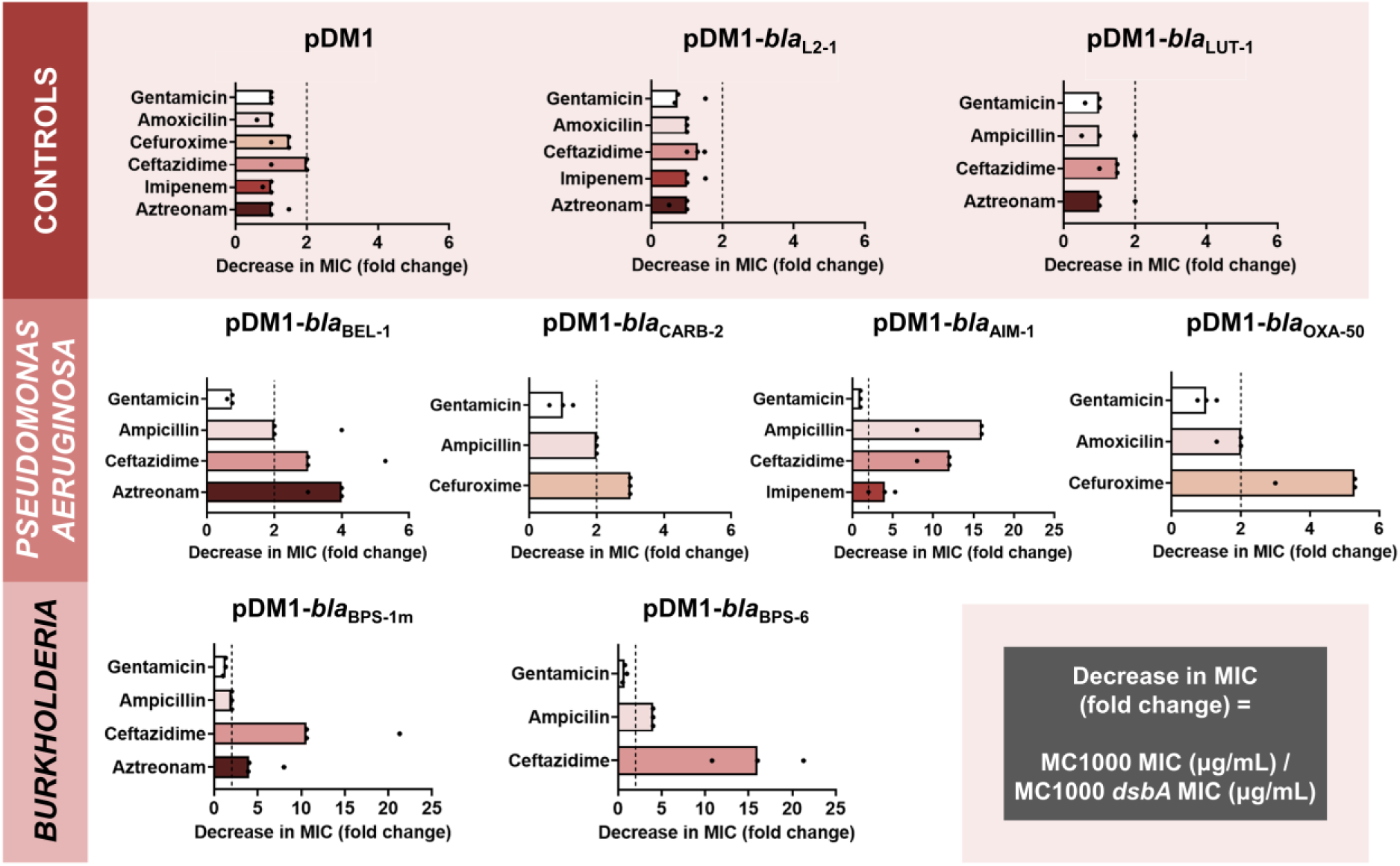
The function of species-specific cysteine-containing β-lactamases from cystic-fibrosis-associated pathogens depends on DsbA-mediated oxidative protein folding. β-lactam MIC values for *E. coli* MC1000 expressing diverse disulfide-bond-containing β-lactamases (Ambler classes A, B and D) are substantially reduced in the absence of DsbA (MIC fold changes: >2; fold change of 2 is indicated by the black dotted lines). No changes in MIC values are observed for the aminoglycoside antibiotic gentamicin (white bars) confirming that absence of DsbA does not compromise the general ability of this strain to resist antibiotic stress. Minor changes in MIC values (≤ 2-fold) are observed for strains harboring the empty vector control (pDM1) or those expressing the class A β-lactamases L2-1 and LUT-1, which contain two or more cysteines (Table S1), but no disulfide bonds (top row). Graphs show MIC fold changes for β-lactamase-expressing *E. coli* MC1000 and its *dsbA* mutant from three biological experiments each conducted as a single technical repeat; the MIC values used to generate this figure are presented in File S2A (rows 2-7 and 9-20).

To ensure that effects shown in Fig. 1 are not due to factors that are not specific to the interaction of DsbA with the tested β-lactamases, we also performed a series of control experiments. We have previously shown that deletion of *dsbA* does not affect the aerobic growth of *E. coli* MC1000, or the permeability of its outer and inner membranes [23]. Furthermore, here we observed no changes in MIC values for the aminoglycoside antibiotic gentamicin, which is not degraded by β-lactamases, or between the parental *E. coli* strain and its *dsbA* mutant harboring only the empty vector (Fig. 1 and File S2A). In addition, *E. coli* strains expressing either of two disulfide-free enzymes, the class A β-lactamases L2-1 and LUT-1 from *S. maltophilia* and *Pseudomonas luteola*, respectively, did not exhibit decreased MICs in the absence of *dsbA* (Fig. 1 and File S2A). These proteins were selected because they both contain two or more cysteine residues, but lack disulfide bonds due to the fact that they are transported to the periplasm, pre-folded, by the Twin-arginine translocation (Tat) pathway, rather than by the Sec system. In the case of L2-1, Tat-dependent transport has been experimentally confirmed [39], whilst LUT-1 contains a predicted Tat signal sequence (SignalP 5.0 [40] likelihood scores: Sec/SPI = 0.0572, Tat/SPI = 0.9312, Sec/SPII (lipoprotein) = 0.0087, other = 0.0029). Finally, the specific interaction between DsbA and our selected test enzymes was further supported by the fact that complementation of *dsbA* generally restores MICs to near wild-type values for the latest generation β-lactam that each β-lactamase can hydrolyze (Fig. S1); we only achieve partial complementation for the *dsbA* mutant expressing BPS-1m, which we attribute to the fact that expression of this enzyme in *E. coli* is sub-optimal.

Taken together, our data show that DsbA-mediated disulfide bond formation is important for the function of all tested, species-specific β-lactamases. Of these, the most affected enzymes (largest MIC value decreases; Fig. 1 and File S2A) are the class A extended-spectrum-β-lactamases (ESBLs) from *Burkholderia* (BPS-1m and BPS-6) and the class B3 metallo-β-lactamase AIM-1, which, like all other class B enzymes [41], is resistant to inhibition by classical β-lactamase inhibitor compounds (Table S1) [30].

#### β-Lactamase abundance and folding

To gain insight into how impairment of disulfide bond formation impacts the production or activity of the tested enzymes (Fig. 1), we first performed immunoblotting for all phylogenetically distinct β-lactamases (AIM-1, BEL-1, OXA-50, CARB-2, and BPS-1m) to assess their protein levels in the presence and absence of *dsbA*. For four of the five tested β-lactamases (AIM-1, BEL-1, OXA-50, and CARB-2) deletion of *dsbA* resulted in drastically reduced protein levels compared to the levels of the control enzyme L2-1, which remained largely unaffected (Fig. 2A). This shows that without their disulfide bonds, these proteins are unstable and are ultimately degraded by other cell envelope proteostasis components [42]. This was further corroborated by the fact that lysates from *dsbA* mutants expressing these four enzymes showed significantly reduced hydrolytic activity towards the chromogenic β-lactamase substrate nitrocefin (Fig. 2B). In the case of BPS-1m, enzyme levels were unchanged in the absence of *dsbA* (Fig. 2A). However, without its disulfide bond, this protein was significantly less able to hydrolyze nitrocefin (Fig. 2B), suggesting a folding defect that results in loss of function. The latter is consistent with the reduced MICs conferred by BPS-1m (and its sister enzyme BPS-6) in the absence of *dsbA* (Fig. 1). The data presented so far (Fig. 1 and 2) demonstrate that disulfide bond formation is essential for the biogenesis (stability and/or protein folding) and, in turn, activity of an expanded set of clinically important β-lactamases, including enzymes that currently lack inhibitor options.

**Figure 2.**
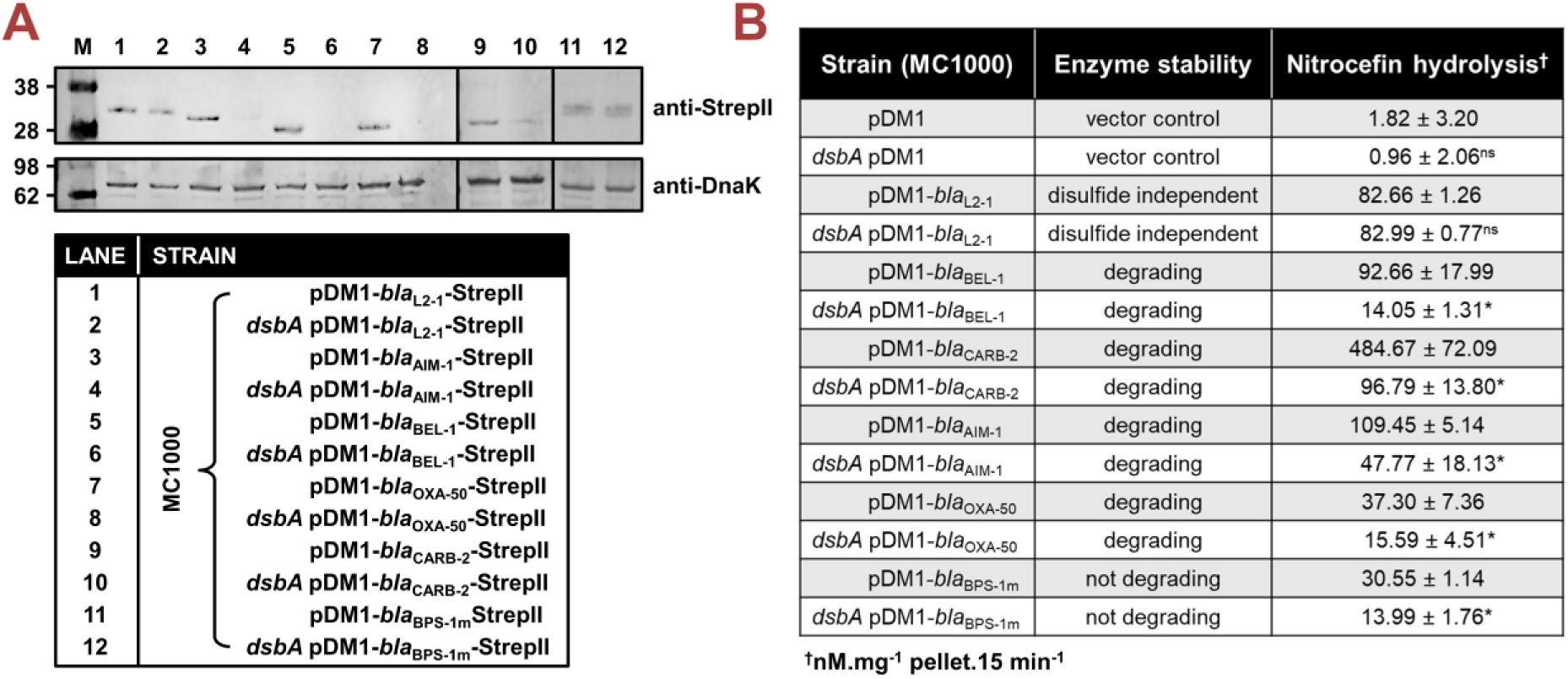
Absence of DsbA results in degradation or misfolding of species-specific cysteine-containing β-lactamases. **(A)** The protein levels of most tested disulfide-bond-containing Ambler class A, B, and D β-lactamases are drastically reduced when these enzymes are expressed in *E. coli* MC1000 *dsbA*; the amount of the control enzyme L2-1, containing three cysteines but no disulfide bonds, is unaffected. An exception to this is the class A enzyme BPS-1m for which no decrease in abundance is observed in the *dsbA* mutant (compare lanes 11 and 12). Protein levels of StrepII-tagged β-lactamases were assessed using a Strep-Tactin-AP conjugate. A representative blot from three biological experiments, each conducted as a single technical repeat, is shown; molecular weight markers (M) are on the left, DnaK was used as a loading control and solid black lines indicate where the membrane was cut. Full immunoblots and SDS PAGE analysis of the immunoblot samples for total protein content are shown in File S3. **(B)** The hydrolysis of the chromogenic β-lactam nitrocefin by cysteine-containing β-lactamases is impaired when these enzymes are expressed in *E. coli* MC1000 *dsbA*. The hydrolytic activities of strains harboring the empty vector or expressing the control enzyme L2-1 show no dependence on DsbA. The “Enzyme stability” column informs on the abundance of each enzyme when it is lacking its disulfide bond(s); this was informed from the immunoblotting experiments in panel (A). The “Nitrocefin hydrolysis” column shows the amount of nitrocefin hydrolyzed per mg of bacterial cell pellet in 15 minutes. n=3, table shows means ±SD, significance is indicated by * = p < 0.05, ns = non-significant.

### Targeting oxidative protein folding inhibits both antibiotic resistance and interbacterial interactions in CF-associated pathogens

#### Sensitization of multidrug-resistant P. aeruginosa clinical isolates

The efficacy of commonly used treatment options against *P. aeruginosa* in CF lung infections, namely piperacillin-tazobactam and cephalosporin-avibactam combinations, as well as more advanced drugs like aztreonam or carbapenems [43,44], is increasingly threatened by an array of β-lactamases, encompassing both broadly disseminated enzymes and species-specific ones [43–45]. To determine whether the effects on β-lactam MICs observed in our inducible system (Fig. 1 and [23]) can be reproduced in the presence of other resistance determinants in a natural context with endogenous enzyme expression levels, we deleted the principal *dsbA* gene, *dsbA1*, in several multidrug-resistant (MDR) *P. aeruginosa* clinical strains (Table S2). Pathogenic bacteria often encode multiple DsbA analogues [27,28] and *P. aeruginosa* is no exception. It encodes two DsbAs, but DsbA1 has been found to catalyze the vast majority of the oxidative protein folding reactions taking place in its cell envelope [46].

We first tested two clinical isolates (strains G4R7 and G6R7; Table S2) expressing the class B3 metallo-β-lactamase AIM-1, for which we recorded reduced activity in an *E. coli dsbA* background (Fig. 1 and 2). This enzyme confers high-level resistance to piperacillin-tazobactam and the third generation cephalosporin ceftazidime, both anti-pseudomonal β-lactams that are used in the treatment of critically ill patients [47]. Notably, while specific to the *P. aeruginosa* genome, *aim-1* is flanked by two ISCR15 elements suggesting that it remains mobilizable [47] (Table S1). MICs for piperacillin-tazobactam and ceftazidime were determined for both AIM-1-positive *P. aeruginosa* isolates and their *dsbA1* mutants (Fig. 3AB). Deletion of *dsbA1* from *P. aeruginosa* G4R7 resulted in a substantial decrease in its piperacillin-tazobactam MIC value by 192 µg/mL and sensitization to ceftazidime (Fig. 3A), while the *dsbA1* mutant of *P. aeruginosa* G6R7 became susceptible to both antibiotic treatments (Fig. 3B). Despite the fact that *P. aeruginosa* G4R7 *dsbA1* was not sensitized for piperacillin-tazobactam, possibly due to the high level of piperacillin-tazobactam resistance of the parent clinical strain, our results across these two isolates show promise for DsbA as a target against β-lactam resistance in *P. aeruginosa*. To further test our approach in an infection context, we performed *in vivo* survival assays using the wax moth model *Galleria mellonella* (Fig. 3C), an informative non-vertebrate system for the study of new antimicrobial approaches against *P. aeruginosa* [48]. Larvae were infected with *P. aeruginosa* G6R7 or its *dsbA1* mutant, and infections were treated once with piperacillin at a final concentration below the EUCAST breakpoint, as appropriate. No larvae survived beyond 20 hours post infection when infected with *P. aeruginosa* G6R7 or its *dsbA1* mutant without antibiotic treatment (Fig. 3C; blue and light blue survival curves). Despite this clinical strain being resistant to piperacillin *in vitro* (Fig. 3B), treatment with piperacillin *in vivo* increased larval survival (52.5% survival at 28 hours post infection) compared to the untreated conditions (Fig. 3C; blue and light blue survival curves) possibly due to *in vivo* ceftazidime MIC values being discrepant to the value recorded *in vitro*. Nonetheless, treatment of *P. aeruginosa* G6R7 *dsbA1* with piperacillin resulted in a significant improvement in survival (77.5% survival at 28 hours post infection), highlighting increased relative susceptibility compared to the treated wild-type condition (Fig. 3C; compare the red and pink survival curves).

**Figure 3.**
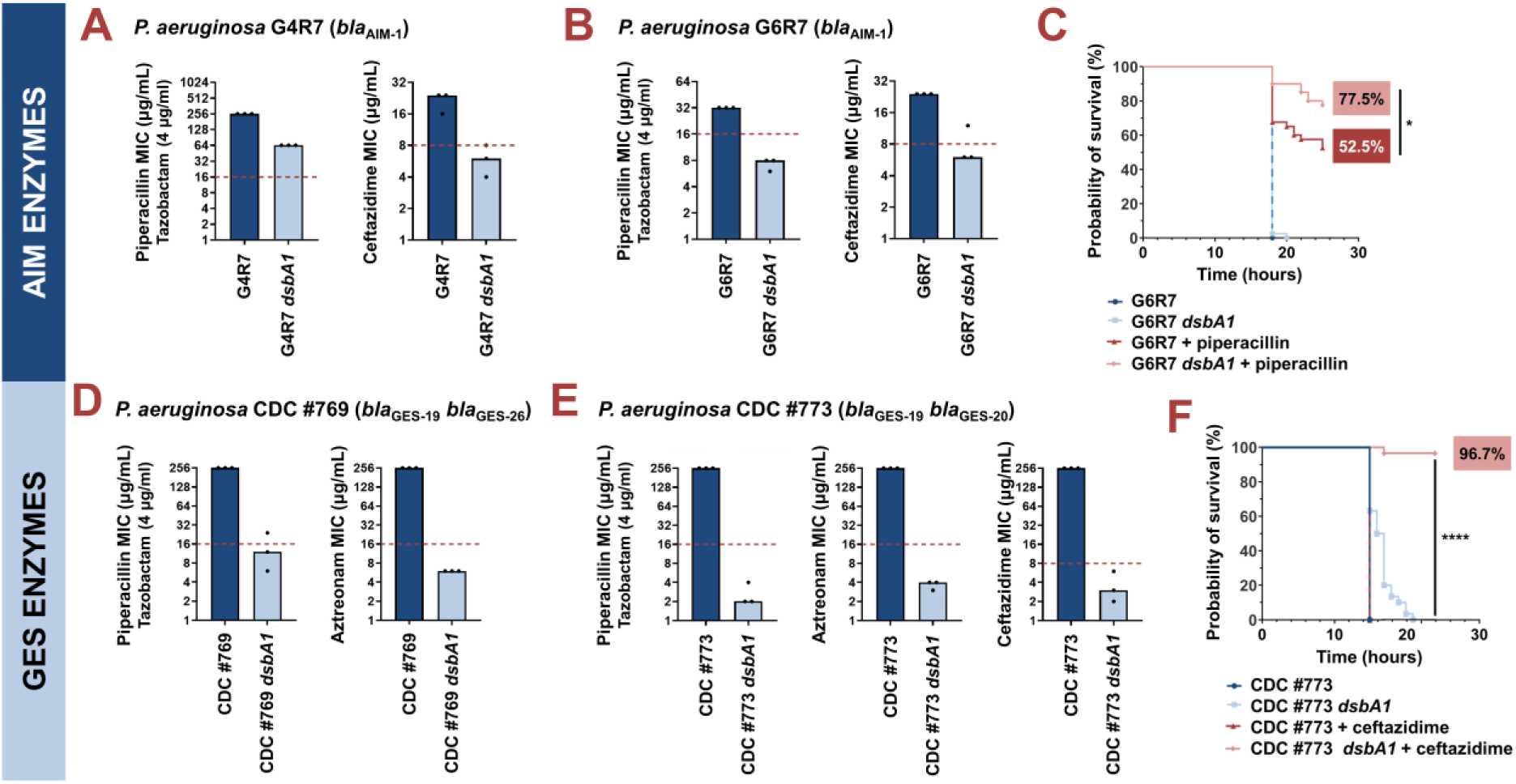
Absence of the principal DsbA analogue (DsbA1) allows treatment of multidrug-resistant *Pseudomonas aeruginosa* clinical isolates with existing β-lactam antibiotics. (A) Deletion of *dsbA1* in the AIM-1-expressing *P. aeruginosa* G4R7 clinical isolate sensitizes this strain to ceftazidime and results in reduction of the piperacillin/tazobactam MIC value by 192 μg/mL. **(B)** Deletion of *dsbA1* in the AIM-1-expressing *P. aeruginosa* G6R7 clinical isolate sensitizes this strain to piperacillin/tazobactam and ceftazidime. **(C)** 100% of the *G. mellonella* larvae infected with *P. aeruginosa* G6R7 (blue curve) or *P. aeruginosa* G6R7 *dsbA1* (light blue curve) die 18 hours post infection, while only 52.5% of larvae infected with *P. aeruginosa* G6R7 and treated with piperacillin (red curve) survive 28 hours post infection. Treatment of larvae infected with *P. aeruginosa* G6R7 *dsbA1* with piperacillin (pink curve) results in 77.5% survival, 28 hours post infection. The graph shows Kaplan-Meier survival curves of infected *G. mellonella* larvae after different treatment applications; horizontal lines represent the percentage of larvae surviving after application of each treatment at the indicated time point (a total of 40 larvae were used for each curve). Statistical analysis of this data was performed using a Mantel-Cox test. The most relevant comparison is noted on the figure. Full statistical analysis is as follows: n=40; p=0.3173 (non-significance) (*P. aeruginosa* vs *P. aeruginosa dsbA1*), p<0.0001 (significance) (*P. aeruginosa* vs *P. aeruginosa* treated with piperacillin), p<0.0001 (significance) (*P. aeruginosa dsbA1* vs *P. aeruginosa* treated with piperacillin), p=0.0147 (significance) (*P. aeruginosa* treated with piperacillin vs *P. aeruginosa dsbA1* treated with piperacillin). **(D)** Deletion of *dsbA1* in the GES-19/GES-26-expressing *P. aeruginosa* CDC #769 clinical isolate sensitizes this strain to piperacillin/tazobactam and aztreonam. **(E)** Deletion of *dsbA1* in the GES-19/GES-20-expressing *P. aeruginosa* CDC #773 clinical isolate sensitizes this strain to piperacillin/tazobactam, aztreonam, and ceftazidime. **(F)** 100% of *G. mellonella* larvae infected with *P. aeruginosa* CDC #773 (blue curve), *P. aeruginosa* CDC #773 *dsbA1* (light blue curve) or larvae infected with *P. aeruginosa* CDC #773 and treated with ceftazidime (red curve) die 21 hours post infection. Treatment of larvae infected with *P. aeruginosa* CDC #773 *dsbA1* with ceftazidime (pink curve) results in 96.7% survival, 24 hours post infection. The graph shows Kaplan-Meier survival curves of infected *G. mellonella* larvae after different treatment applications; horizontal lines represent the percentage of larvae surviving after application of each treatment at the indicated time point (a total of 30 larvae were used for each curve). Statistical analysis of this data was performed using a Mantel-Cox test. The most relevant comparison is noted on the figure. Full statistical analysis is as follows: n=30; p<0.0001 (significance) (*P. aeruginosa* vs *P. aeruginosa dsbA1*), p>0.9999 (non-significance) (*P. aeruginosa* vs *P. aeruginosa* treated with ceftazidime), p<0.0001 (significance) (*P. aeruginosa dsbA1* vs *P. aeruginosa* treated with ceftazidime), p<0.0001 (significance) (*P. aeruginosa* treated with ceftazidime vs *P. aeruginosa dsbA1* treated with ceftazidime). For panels (A), (B), (D), and (E) the graphs show MIC values (μg/mL) from three biological experiments, each conducted as a single technical repeat; red dotted lines indicate the EUCAST clinical breakpoint for each antibiotic.

Next, we tested two *P. aeruginosa* clinical isolates (strains CDC #769 and CDC #773; Table S2), each expressing two class A enzymes from the GES family (GES-19/GES-26 or GES-19/GES-20), for which we have previously demonstrated DsbA dependence [23]. The GES family comprises 59 distinct ESBLs (File S1), which are globally disseminated and commonly found in *P. aeruginosa*, as well as other critical Gram-negative pathogens (for example *Klebsiella pneumoniae* and *Enterobacter cloacae*) [49]. Deletion of *dsbA1* in these clinical strains resulted in sensitization to piperacillin-tazobactam and aztreonam for *P. aeruginosa* CDC #769 (Fig. 3D), and to representative compounds of all classes of anti-pseudomonal β-lactam drugs (piperacillin-tazobactam, aztreonam, and ceftazidime) for *P. aeruginosa* CDC #773 (Fig. 3E). *P. aeruginosa* CDC #773 and its *dsbA1* mutant were further tested in a *G. mellonella* infection model using ceftazidime treatment (Fig. 3F). In this case, no larvae survived 24 hours post infection (Fig. 3F; blue, light blue and red survival curves), except for insects infected with *P. aeruginosa* CDC #773 *dsbA1* and treated with ceftazidime at a final concentration below the EUCAST breakpoint, whereby 96.7% survival was recorded (Fig. 3F; pink survival curves).

We have demonstrated the specific interaction of DsbA with the tested β-lactamase enzymes in our *E. coli* K-12 inducible system using gentamicin controls (Fig. 1 and File S2A) and gene complementation (Fig. S1). To confirm the specificity of this interaction in *P. aeruginosa*, we performed representative control experiments in one of our clinical strains, *P. aeruginosa* CDC #769. We first tested the general ability of *P. aeruginosa* CDC #769 *dsbA1* to resist antibiotic stress by recording MIC values against gentamicin, and found it unchanged compared to its parent (Fig. S2A). Gene complementation in clinical isolates is especially challenging and rarely attempted due to the high levels of resistance and lack of genetic tractability in these strains. Despite these challenges, to further ensure the specificity of the interaction of DsbA with tested β-lactamases in *P. aeruginosa*, we have complemented *dsbA1* from *P. aeruginosa* PAO1 into *P. aeruginosa* CDC #769 *dsbA1*. We found that complementation of *dsbA1* restores MICs to wild-type values for both tested β-lactam compounds (Fig. S2B) further demonstrating that our results in *P. aeruginosa* clinical strains are not confounded by off-target effects.

Our data on the sensitization of AIM– and GES-expressing *P. aeruginosa* clinical isolates to commonly used anti-pseudomonal β-lactam drugs, combined with our previous results on strains producing β-lactamases from the OXA family [23], show that our approach holds promise towards inactivating numerous clinically important *Pseudomonas*-specific enzymes. These include resistance determinants that cannot be currently targeted by classical β-lactamase inhibitor compounds (for example enzymes from the OXA and AIM families [30]) and, therefore, limit our treatment options.

#### New treatment options for extremely-drug-resistant S. maltophilia clinical isolates

We have previously used our inducible *E. coli* K-12 experimental system to demonstrate that the function of the inhibitor-resistant class B3 metallo-β-lactamase L1-1 from *S. maltophilia* is dependent on DsbA [23]. By contrast, the second β-lactamase encoded on the chromosome of this species, L2-1, which we use as a negative control in this study (Fig. 1 and 2), is not DsbA dependent. The hydrolytic spectra of these β-lactamases are exquisitely complementary [13,14], making this bacterium resistant to most β-lactam compounds commonly used for CF patients. Considering that L1 enzymes are the sole drivers of ceftazidime resistance, we wanted to investigate the DsbA dependency of L1-1 in its natural context to determine whether inhibition of oxidative protein folding potentiates the activity of complex cephalosporins against this pathogen.

We compromised disulfide bond formation in two clinical isolates of *S. maltophilia* (strains AMM and GUE; Table S2), by deleting the main *dsbA* gene cluster (directly adjacent *dsbA* and *dsbL* genes, with DsbL predicted to be a DsbA analogue [28]) and recorded a drastic decrease of ceftazidime MIC values for both mutant strains (Fig. 4A,B). Since *S. maltophilia* cannot be treated with ceftazidime, there is no EUCAST breakpoint available for this organism. That said, for both tested *dsbA dsbL* mutant strains, the recorded ceftazidime MIC values were lower than the ceftazidime EUCAST breakpoint for the related major pathogen *P. aeruginosa* [50].

**Figure 4.**
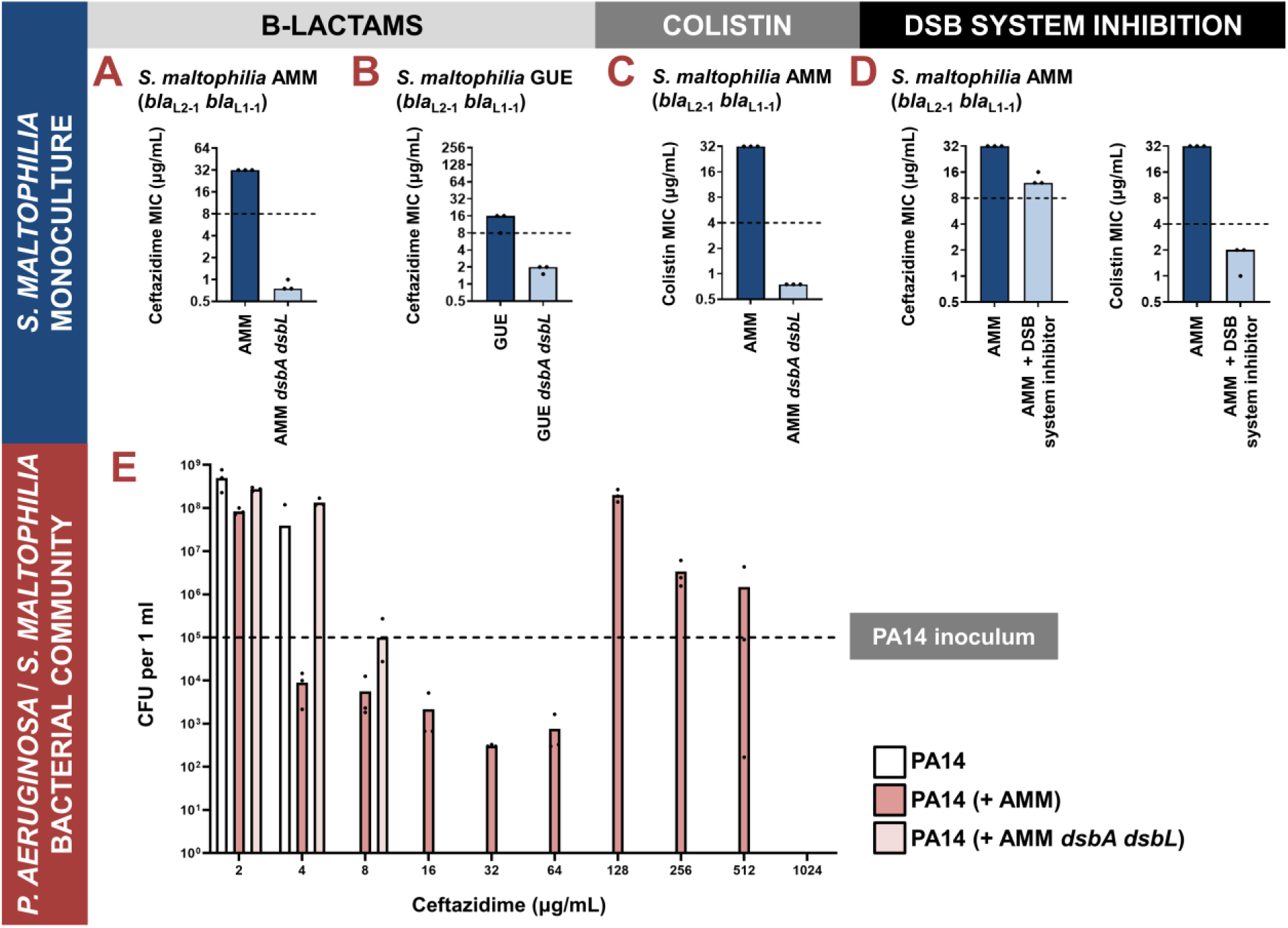
(A-D) Impairment of disulfide bond formation allows the treatment of *Stenotrophomonas maltophilia* clinical strains with β-lactam and colistin antibiotics. (A, B) Deletion of *dsbA dsbL* in the *S. maltophilia* AMM and *S. maltophilia* GUE clinical isolates results in drastic decrease of their ceftazidime MIC values. (C) Deletion of *dsbA dsbL* in the *S. maltophilia* AMM clinical strain results in drastic decrease of its colistin MIC value. (D) Use of a small-molecule inhibitor of DsbB against the *S. maltophilia* AMM clinical strain results in decrease of its ceftazidime and colistin MIC values. For panels (A-D) graphs show MIC values (μg/mL) from three biological experiments; for β-lactam MIC assays each experiment was conducted as a single technical repeat, whereas for colistin MIC assays each experiment was conducted in technical triplicate. In the absence of EUCAST clinical breakpoints for *S. maltophilia*, the black dotted lines indicate the EUCAST clinical breakpoint for each antibiotic for the related pathogen *P. aeruginosa*. (E) Protection of *P. aeruginosa* by *S. maltophilia* clinical strains is dependent on oxidative protein folding. The susceptible *P. aeruginosa* strain PA14 can survive exposure to ceftazidime up to a maximum concentration of 4 μg/mL when cultured in isolation (white bars). By contrast, if co-cultured in the presence of *S. maltophilia* AMM, which can hydrolyze ceftazidime through the action of its L1-1 β-lactamase enzyme, *P. aeruginosa* PA14 can survive and actively grow in concentrations of ceftazidime as high as 512 μg/mL (dark pink bars). This protection is abolished if *P. aeruginosa* PA14 is co-cultured with *S. maltophilia* AMM *dsbA dsbL* (light pink bars), where L1-1 is inactive (as shown in Fig. 4A and [23]). The graph shows *P. aeruginosa* PA14 colony forming unit counts (CFUs) for each condition; three biological replicates were conducted in technical triplicate, and mean CFU values are shown. The black dotted line indicates the *P. aeruginosa* PA14 inoculum. The mean CFU values used to generate this figure are presented in File S2B.

In addition to being resistant to β-lactams, *S. maltophilia* is usually intrinsically resistant to colistin [12], which precludes the use of yet another broad class of antibiotics. Bioinformatic analysis on 106 complete *Stenotrophomonas* genomes revealed that most strains of this organism carry two chromosomally-encoded MCR analogues that cluster with clinical MCR-5 and MCR-8 proteins (File S4). We have previously found the activity of all clinical MCR enzymes to be dependent on the presence of DsbA [23], thus we compared the colistin MIC value of the *S. maltophilia* AMM *dsbA dsbL* strain to that of its parent. We found that impairment of disulfide bond formation in this strain resulted in a decrease of its colistin MIC value from 32 μg/mL to 0.75 μg/mL (Fig. 4C). Once more, there is no colistin EUCAST breakpoint available for *S. maltophilia*, but a comparison with the colistin breakpoint for *P. aeruginosa* (4 μg/mL) demonstrates the magnitude of the effects that we observe.

Since the *dsbA* and *dsbL* are organized in a gene cluster in *S. maltophilia*, we wanted to ensure that our results reported above were exclusively due to disruption of disulfide bond formation in this organism. First, we recorded gentamicin MIC values for *S. maltophilia* AMM *dsbA dsbL* and found them to be unchanged compared to the gentamicin MICs of the parent strain (Fig. S2C). This confirms that disruption of disulfide bond formation does not compromise the general ability of this organism to resist antibiotic stress. Next, we complemented *S. maltophilia* AMM *dsbA dsbL*. The specific oxidative roles and exact regulation of DsbA and DsbL in *S. maltophilia* remain unknown. For this reason and considering that genetic manipulation of extremely-drug-resistant organisms is challenging, we used our genetic construct optimized for complementing *P. aeruginosa* CDC #769 *dsbA1* with *dsbA1* from *P. aeruginosa* PAO1 (Fig. S2B) to also complement *S. maltophilia* AMM *dsbA dsbL*. We based this approach on the fact that DsbA proteins from one species have been commonly shown to be functional in other species [51–54]. Indeed, we found that complementation of *S. maltophilia* AMM *dsbA dsbL* with *P. aeruginosa* PAO1 *dsbA1* restores MICs to wild-type values for both ceftazidime and colistin (Fig. S2D), conclusively demonstrating that our results in *S. maltophilia* are not confounded by off-target effects.

The DSB proteins have been shown to play a central role in bacterial virulence, and in this context, they have been proposed as promising targets against bacterial pathogenesis [27,28,55]. As a result, several laboratory compounds against both DsbA [56,57] and its partner protein DsbB [35], which maintains DsbA in a catalytically active state [58], have been developed. We have successfully used one of these inhibitors, 4,5-dichloro-2-(2-chlorobenzyl)pyridazin-3-one, termed “compound 12” in (47), to achieve sensitization of clinical strains of Enterobacteria to β-lactam and colistin antibiotics [23]. Here, we used a derivative compound, 4,5-dibromo-2-(2-chlorobenzyl)pyridazin-3(2H)-one, termed “compound 36” in [59], which is an improved analog of compound 12 and has been shown to target several DsbB proteins from Gram-negative pathogens that share 20-80% in protein identity. Compound 36 was previously shown to inhibit disulfide bond formation in *P. aeruginosa* via covalently binding onto one of the four essential cysteine residues of DsbB in the DsbA-DsbB complex [59]. Since *S. maltophilia* DsbB shares ∼28% protein sequence identity with analogues from *P. aeruginosa*, we reasoned that this pathogen could be a good candidate for testing DSB system inhibition. Exposure of *S. maltophilia* AMM to the DSB inhibitor lowered its ceftazidime MIC value by at least 16-20 μg/mL and decreased its colistin MIC value from 32 μg/mL to 2 μg/mL (Fig. 4D); this decrease in the colistin MIC is commensurate with the results we obtained for the *S. maltophilia* AMM *dsbA dsbL* strain (Fig. 4C). The activity of compound 36 is specific to inhibition of disulfide bond formation since the gentamicin MIC values of *S. maltophilia* AMM remain unchanged in the presence of the inhibitor and treatment of *S. maltophilia* AMM *dsbA dsbL* with the compound does not affect its colistin MIC value (Fig. S2E). Considering that this inhibitor has not been specifically optimized for *S. maltophilia* strains, the recorded drops in MIC values (Fig. 4D) are encouraging and suggest that the DSB system proteins are tractable targets against species-specific resistance determinants in this pathogen.

Currently, the best clinical strategy against *S. maltophilia* is to reduce the likelihood of infection [60], therefore novel treatment strategies against this organism are desperately needed. Overall, our results on targeting oxidative protein folding in this organism show promise for the generation of therapeutic avenues that are compatible with mainstream antibiotics (β-lactams and polymyxins), which are commonly used for the treatment of other pathogens, for example *P. aeruginosa*, in CF lung infections.

#### Inhibition of cross-resistance in S. maltophilia – P. aeruginosa mixed communities

The antibiotic resistance mechanisms of *S. maltophilia* impact the antibiotic tolerance profiles of other organisms that are found in the same infection environment. *S. maltophilia* hydrolyses all β-lactam drugs through the action of its L1 and L2 β-lactamases [13,14]. In doing so, it has been experimentally shown to protect other pathogens that are, in principle, susceptible to treatment, such as *P. aeruginosa [15*]. This protection, in turn, allows active growth of otherwise treatable *P. aeruginosa* in the presence of complex β-lactams, like imipenem [15], and, at least in some conditions, increases the rate of resistance evolution of *P. aeruginosa* against these antibiotics [17].

We wanted to investigate whether our approach would be useful in abrogating interspecies interactions that are relevant to CF infections. We posited that ceftazidime resistance in *S. maltophilia* is largely driven by L1-1, an enzyme that we can incapacitate by targeting disulfide bond formation [23] (Fig. 4A,B,D). As such, impairment of oxidative protein folding in *S. maltophilia* should allow treatment of this organism with ceftazidime, and at the same time eliminate any protective effects that benefit susceptible strains of co-occurring organisms. With ceftazidime being a standard anti-pseudomonal drug, and in view of the interactions reported between *P. aeruginosa* and *S. maltophilia* [15,17,61], we chose to test this hypothesis using *S. maltophilia* AMM and a *P. aeruginosa* strain that is sensitive to β-lactam antibiotics, *P. aeruginosa* PA14. We followed established co-culture protocols for these organisms [15] and first monitored the survival and growth of *P. aeruginosa* under ceftazidime pressure in monoculture, or in the presence of *S. maltophilia* strains. Due to the naturally different growth rates of these two species (*S. maltophilia* grows much slower than *P. aeruginosa*) especially in laboratory conditions, the protocol we followed [15] requires *S. maltophilia* to be grown for 6 hours prior to co-culturing it with *P. aeruginosa*. To ensure that at this point in the experiment our two *S. maltophilia* strains, with and without *dsbA*, had grown comparatively to each other, we determined their cell densities (Fig. S3A). We found that *S. maltophilia* AMM *dsbA dsbL* had grown at a similar level as the wild-type strain, and both were at a higher cell density [∼10^7^ colony forming units (CFUs)] compared to the *P. aeruginosa* PA14 inoculum (5 x 10^4^ CFUs).

*P. aeruginosa* PA14 monoculture cannot grow in the presence of more than 4 μg/mL of ceftazidime (Fig. 4E; white bars). However, the same strain can actively grow in concentrations of ceftazidime up to 512 μg/mL in the presence of *S. maltophilia* AMM (Fig. 4E; dark pink bars), showing that the protective effects previously observed with imipenem [15] are applicable to other clinically relevant β-lactam antibiotics. Cross-resistance effects are most striking at concentrations of ceftazidime above 64 μg/ml; for amounts between 16 and 64 μg/ml, *P. aeruginosa* survives in the presence of *S. maltophilia*, but does not actively grow. This is in agreement with previous observations showing that the expression of L1-1 is induced by the presence of complex β-lactams [62]. In this case, the likely increased expression of L1-1 in *S. maltophilia* grown in concentrations of ceftazidime equal or higher than 128 μg/ml promotes ceftazidime hydrolysis and decrease of the active antibiotic concentration, in turn, shielding the susceptible *P. aeruginosa* strain. By contrast, protective effects are almost entirely absent when *P. aeruginosa* PA14 is co-cultured with *S. maltophilia* AMM *dsbA dsbL*, which cannot hydrolyze ceftazidime efficiently because L1-1 activity is impaired [23] (Fig. 4A,B,D). In fact, in these conditions *P. aeruginosa* PA14 only survives in concentrations of ceftazidime up to 8 μg/mL (Fig. 4E; light pink bars), 64-fold lower than what it can endure in the presence of *S. maltophilia* AMM (Fig. 4E; dark pink bars).

To ensure that ceftazidime treatment leads to eradication of both *P. aeruginosa* and *S. maltophilia* when disulfide bond formation is impaired in *S. maltophilia*, we monitored the abundance of both strains in each synthetic community for select antibiotic concentrations (Fig. S3B). In this experiment we largely observed the same trends as in Fig. 4E. At low antibiotic concentrations, for example 4 μg/mL of ceftazidime, *S. maltophilia* AMM is fully resistant and thrives, thus outcompeting *P. aeruginosa* PA14 (dark pink and dark blue bars in Fig. S3B). The same can also be seen in Fig. 4E, whereby decreased *P. aeruginosa* PA14 CFUs are recorded. By contrast *S. maltophilia* AMM *dsbA dsbL* already displays decreased growth at 4 μg/mL of ceftazidime because of its non-functional L1-1 enzyme, allowing comparatively higher growth of *P. aeruginosa* (light pink and light blue bars in Fig. S3B). Despite the competition between the two strains, *P. aeruginosa* PA14 benefits from *S. maltophilia* AMM’s high hydrolytic activity against ceftazidime, which allows it to survive and grow in high antibiotic concentrations even though it is not resistant (see 128 μg/mL; dark pink and dark blue bars in Fig. S3B). In stark opposition, without its disulfide bond in *S. maltophilia* AMM *dsbA dsbL*, L1-1 cannot confer resistance to ceftazidime, resulting in killing of *S. maltophilia* AMM *dsbA dsbL* and, consequently, also of *P. aeruginosa* PA14 (see 128 μg/mL; light pink and light blue bars in Fig. S3B).

The data presented here show that, at least under laboratory conditions, targeting protein homeostasis pathways in specific recalcitrant pathogens has the potential to not only alter their own antibiotic resistance profiles (Fig. 3 and 4A-D), but also to influence the antibiotic susceptibility profiles of other bacteria that co-occur in the same conditions (Fig. 5). Admittedly, the conditions in a living host are too complex to draw direct conclusions from this experiment. That said, our results show promise for infections, where pathogen interactions affect treatment outcomes, and whereby their inhibition might facilitate treatment.

**Figure 5.**
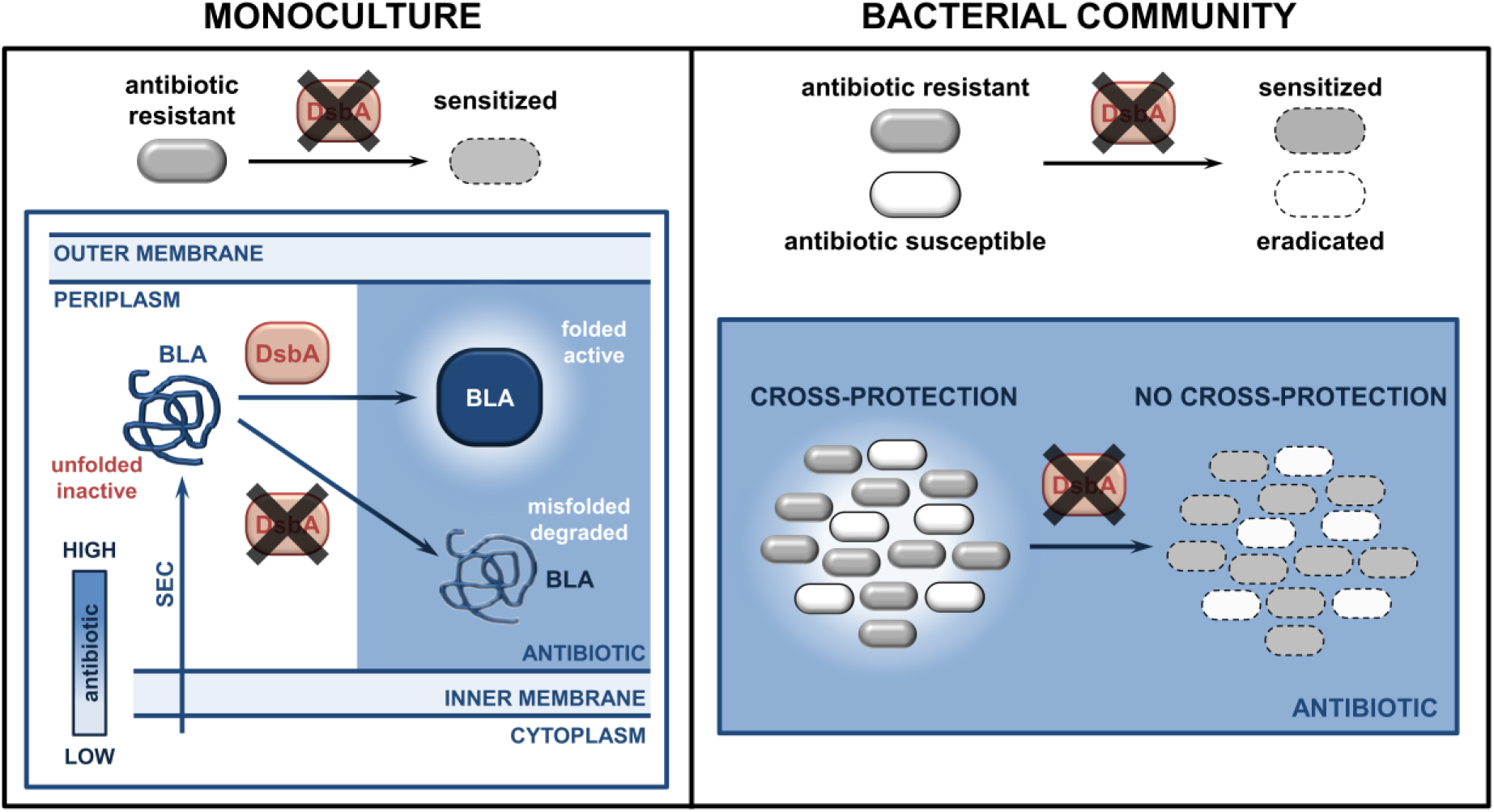
Inhibition of oxidative protein folding counters antibiotic resistance and inter-species interactions in CF-associated pathogens. (Left) After Sec translocation to the periplasm and DsbA-assisted folding, cysteine-containing species-specific β-lactamase enzymes from recalcitrant pathogens, like *P. aeruginosa* or *S. maltophilia*, are active and can hydrolyze β-lactam antibiotics. However, in the absence of their disulfide bonds, DsbA-dependent β-lactamases either degrade or misfold, and thus can no longer confer resistance to β-lactam compounds. (Right) In multispecies bacterial communities, bacteria that degrade antibiotics, for example species producing β-lactamases, can protect antibiotic-susceptible strains. Targeting disulfide bond formation impairs interbacterial interactions that are reliant on the activity of DsbA-dependent β-lactamase enzymes, allowing eradication of both bacterial species.

## DISCUSSION

Impairment of cell envelope protein homeostasis through interruption of disulfide bond formation has potential as a broad-acting strategy against AMR in Gram-negative bacteria [23]. Here, we focus on the benefits of such an approach against pathogens encountered in challenging infection settings by studying organisms found in the CF lung. In particular, we show that incapacitation of oxidative protein folding compromises the function of diverse β-lactamases that are specific to CF-associated bacteria, like *P. aeruginosa* and *Burkholderia* complex (Fig. 1 and 2). Furthermore, we find that the effects we observe at the enzyme level are applicable to multiple MDR *P. aeruginosa* and extremely-drug-resistant *S. maltophilia* clinical strains, both *in vitro* (Fig. 3A,B,D,E and 4A,B) and in an *in vivo* model of infection (Fig. 3C,F). Our findings, so far, concern β-lactamases encoded by enteric pathogens (discussed in [23]) or CF-associated organisms (discussed in [23] and in this study). Nonetheless, many other environmental bacteria are opportunistic human pathogens and encode β-lactamase genes that make them highly resistant to antibiotic treatment [13,63,64]. The ubiquitous nature of disulfide bond formation systems across Gram-negative species guarantees that the same approach can be expanded. To provide some proof on this front, we investigated two additional class B3 metallo-β-lactamases, POM-1 produced by *Pseudomonas otitidis* and SMB-1 encoded on the chromosome of *Serratia marcescens* (Table S1). We tested these enzymes in our inducible *E. coli* K-12 system and found that their activities are indeed DsbA dependent (Fig. S4A and S5 and File S2A), with SMB-1 degrading in the absence of DsbA and POM-1 suffering a folding defect (Fig. S4B,C). Since 57% of β-lactamase phylogenetic families that are found in pathogens and organisms capable of causing opportunistic infections contain members with two or more cysteines (File S1), we expect that thousands of enzymes rely on DsbA for their stability and function. Focusing solely on the β-lactamase families that we have investigated here and previously [23] (17 phylogenetic families), we estimate that upwards of 575 discrete proteins are DsbA dependent. This encompasses enzymes specific to pathogens with very limited treatment options, for example the *Burkholderia* complex (Fig. 1 and File S2A) and *S. maltophilia* (Fig. 4A,B,D), as well as 145 β-lactamases that cannot be inhibited by classical adjuvant approaches, like class B enzymes [30] from the AIM, L1, POM, and SMB families (Fig. 1, 3A-C and S4 and File S2A).

Of the organisms studied in this work, *S. maltophilia* deserves further discussion because of its unique intrinsic resistance profile. The prognosis of CF patients with *S. maltophilia* lung carriage is still debated [8,65–72], largely because studies with extensive and well-controlled patient cohorts are lacking. This notwithstanding, the therapeutic options against this pathogen are currently limited to one non-β-lactam antibiotic-adjuvant combination, which is not always effective, trimethoprim-sulfamethoxazole [73–76], and a few last-line β-lactam drugs, like the fifth-generation cephalosporin cefiderocol and the combination aztreonam-avibactam. Resistance to commonly used antibiotics causes many problems during treatment and, as a result, infections that harbor *S. maltophilia* have high case fatality rates [13]. This is not limited to CF patients, as *S. maltophilia* is a major cause of death in children with bacteremia [11]. We find that targeting disulfide bond formation in this species allows its treatment with cephalosporins, like ceftazidime, (Fig. 4A,B,D) and, at the same time, leads to colistin potentiation (Fig. 4C,D). Our results create a foundation for extending the usability of two invaluable broad-acting antibiotic classes against this challenging organism. At the same time, *S. maltophilia* is often found to co-exist in the CF lung with other pathogens like *P. aeruginosa* [8–10]. Even though current studies are confined to laboratory settings [15], it is likely that interactions between these two species makes treatment of polymicrobial infections more complex. Here, we demonstrate that by compromising L1-1 through impairing protein homeostasis in *S. maltophilia* (Fig. 4A,B,D and [23]), in addition to generating new treatment options (Fig. 4A-D), we abolish the capacity of this organism to protect other species (Fig. 4E). Since similar bacterial interactions are documented in resistant infections [6], it can be expected that our approach will yield analogous results for other coexisting CF lung pathogens that produce DsbA-dependent β-lactamases [23], for example *P. aeruginosa* and *S. aureus* [77,78] or *K. pneumoniae* and *Acinetobacter baumannii* [16] (Fig. 5).

More generally, our findings serve as proof of principle of the added benefits of strategies that aim to incapacitate resistance determinants like β-lactamases. These proteins threaten the most widely prescribed class of antibiotics worldwide [79] and, at the same time, can promote cross-resistance between pathogens found in polymicrobial infections. It is therefore important to continue developing β-lactamase inhibitors, which, so far, have been one of the biggest successes in our battle against AMR [30,80]. That said, the deployment of broad-acting small molecules with the capacity to bind and effectively inhibit thousands of clinically important β-lactamases [7741 distinct documented enzymes [81] (File S1)] is challenging and, eventually, leads to the emergence of β-lactamase variants that are resistant to combination therapy. As such, development of additional alternative strategies that can broadly incapacitate these resistance proteins, ideally without the need to bind to their active sites, is critical. This has been shown to be possible through metal chelation for class B metallo-β-lactamases [82]. Adding to this, our previous work [23] and the results presented here lay the groundwork for exploiting accessible cell envelope proteostasis processes to generate new resistance breakers. Inhibiting such systems has untapped potential for the design of broad-acting next-generation therapeutics, which simultaneously compromise multiple resistance mechanisms [23], and also for the development of species-or infection-specific approaches that are well suited for the treatment of complex polymicrobial communities (Fig. 5).

## MATERIALS AND METHODS

### Reagents and bacterial growth conditions

Unless otherwise stated, chemicals and reagents were acquired from Sigma Aldrich or Fisher Scientific, growth media were purchased from Oxoid and antibiotics were obtained from Melford Laboratories. Lysogeny broth (LB) (10 g/L NaCl) and agar (1.5% w/v) were used for routine growth of all organisms at 37 °C with shaking at 220 RPM, as appropriate. Mueller-Hinton (MH) broth and agar (1.5% w/v) were used for Minimum Inhibitory Concentration (MIC) assays. Growth media were supplemented with the following, as required: 0.25 mM Isopropyl β-D-1-thiogalactopyranoside (IPTG), 50 μg/mL kanamycin, 100 μg/mL ampicillin, 33 μg/mL chloramphenicol, 33 μg/mL gentamicin (for cloning purposes), 400-600 μg/mL gentamicin (for genetic manipulation of *P. aeruginosa* and *S. maltophilia* clinical isolates), 12.5 μg/mL tetracycline (for cloning purposes), 100-400 μg/mL tetracycline (for genetic manipulation of *P. aeruginosa* clinical isolates), 50 μg/mL streptomycin (for cloning purposes), 2000-5000 μg/mL streptomycin (for genetic manipulation of *P. aeruginosa* clinical isolates), and 6000 μg/mL streptomycin (for genetic manipulation of *S. maltophilia* clinical isolates).

### Construction of plasmids and bacterial strains

Bacterial strains, plasmids and oligonucleotides used in this study are listed in Tables S2, S3 and S4, respectively. DNA manipulations were conducted using standard methods. KOD Hot Start DNA polymerase (Merck) was used for all PCR reactions according to the manufacturer’s instructions, oligonucleotides were synthesized by Sigma Aldrich and restriction enzymes were purchased from New England Biolabs. All constructs were DNA sequenced and confirmed to be correct before use.

Genes for β-lactamase enzymes were amplified from genomic DNA extracted from clinical isolates (Table S5) with the exception of *bps-1m, bps-6, carb-2, ftu-1* and *smb-1*, which were synthesized by GeneArt Gene Synthesis (ThermoFisher Scientific). β-lactamase genes were cloned into the IPTG-inducible plasmid pDM1 using primers P1-P16. All StrepII-tag fusions of β-lactamase enzymes (constructed using primers P3, P5, P7, P9, P11, P13, P15, and P17-23) have a C-terminal StrepII tag (GSAWSHPQFEK).

*P. aeruginosa dsbA1* mutants and *S. maltophilia dsbA dsbL* mutants were constructed by allelic exchange, as previously described [83]. Briefly, the *dsbA1* gene area of *P. aeruginosa* strains (including the *dsbA1* gene and ∼600 bp on either side of this gene) was amplified (primers P24/P25) and the obtained DNA was sequenced to allow for accurate primer design for the ensuing cloning step. The pKNG101-*dsbA1* plasmid was then used for deletion of the *dsbA1* gene in *P. aeruginosa* G4R7 and *P. aeruginosa* G4R7, as before [23]. For the deletion of *dsbA1* in *P. aeruginosa* CDC #769 and *P. aeruginosa* CDC #773, ∼500-bp DNA fragments upstream and downstream of the *dsbA1* gene were amplified using *P. aeruginosa* CDC #769 or *P. aeruginosa* CDC #773 genomic DNA [primers P28/P29 (upstream) and P30/P31 (downstream)]. Fragments containing both regions were then obtained by overlapping PCR (primers P28/P31) and inserted into the XbaI/BamHI sites of pKNG102, resulting in plasmids pKNG102-*dsbA1*-769 and pKNG102-*dsbA1*-773. For *S. maltophilia* strains the *dsbA dsbL* gene area (including the *dsbA dsbL* genes and ∼1000 bp on either side of these genes) was amplified (primers P26/P27) and the obtained DNA was sequenced to allow for accurate primer design for the ensuing cloning step. Subsequently, ∼700-bp DNA fragments upstream and downstream of the *dsbA dsbL* genes were amplified using *S. maltophilia* AMM or *S. maltophilia* GUE genomic DNA [primers P32/P33 (upstream) and P34/P35 (downstream)]. Fragments containing both of these regions were then obtained by overlapping PCR (primers P32/35) and inserted into the XbaI/BamHI sites of pKNG101, resulting in plasmids pKNG101-*dsbA dsbL*-AMM and pKNG101-*dsbA dsbL*-GUE. The suicide vector pKNG101 [84] and its derivative pKNG102, are not replicative in *P. aeruginosa* or *S. maltophilia*; both vectors are maintained in *E. coli* CC118λpir and mobilized into *P. aeruginosa* and *S. maltophilia* strains by triparental conjugation. For *P. aeruginosa,* integrants were selected on Vogel Bonner Minimal medium supplemented with streptomycin (for *P. aeruginosa* G4R7 and *P. aeruginosa* G6R7) or tetracycline (for *P. aeruginosa* CDC #769 and *P. aeruginosa* CDC #773). For *S. maltophilia,* integrants were selected on MH agar supplemented with streptomycin and ampicillin. Successful integrants were confirmed using PCR, and mutants were resolved by exposure to 20% sucrose. Gene deletions were confirmed via colony PCR and DNA sequencing (primers P24/P25).

*P. aeruginosa* PA14, *S. maltophilia* AMM, and *S. maltophilia* AMM *dsbA dsbL* were labelled with a gentamicin resistance marker using mini-Tn*7* delivery transposon-based vectors adapted from Zobel et al. [85]. The non-replicative vectors pTn7-M (labelling with gentamicin resistance only, for *P. aeruginosa* PA14) and pBG42 (labelling with gentamicin resistance and msfGFP, for *S. maltophilia* strains) were mobilized into the respective recipients using conjugation, in the presence of a pTNS2 plasmid expressing the TnsABC+D specific transposition pathway. Correct insertion of the transposon into the *att*Tn*7* site was confirmed via colony PCR and DNA sequencing (primers P44/P45 for *P. aeruginosa*, primers P46/P47 for *S. maltophilia*).

*P. aeruginosa* CDC #769 *dsbA1* and *S. maltophilia* AMM *dsbA dsbL* were complemented with DsbA1 from *P. aeruginosa* PAO1 using a mini-Tn*7* delivery transposon-based vector adapted from Zobel et al. [85]. Briefly, the *msfGFP* gene of pBG42 was replaced with the *dsbA1* gene of *P. aeruginosa* PAO1 by HiFi DNA assembly according to the manufacturer’s instructions (NEBuilder HiFi DNA Assembly, New England Biolabs). The *dsbA1* gene of *P. aeruginosa* PAO1 was amplified from genomic DNA using primers P38/P39 and the vector was linearized with primers P36/P37. *msfGFP* amplified from pBG42 with primers P40/P41 was reintroduced onto the vector under the PEM7 promoter between the HindIII and BamHI sites of pBG42 [86] resulting in plasmid pBG42-PAO1*dsbA1*. Correct assembly of pBG42-PAO1*dsbA1* was confirmed by colony PCR (primers P42/P43) and DNA sequencing. pBG42-PAO1*dsbA1* was mobilized into the recipient strains using conjugation, in the presence of a pTNS2 plasmid expressing the TnsABC+D specific transposition pathway. GFP positive colonies were screened using colony PCR and correct insertion of the transposon into the *att*Tn*7* site of clinical strains was confirmed via DNA sequencing (primers P44/P45 for *P. aeruginosa*, primers P46/P47 for *S. maltophilia*).

#### Minimum inhibitory concentration (MIC) assays

Unless otherwise stated, antibiotic MIC assays were carried out in accordance with the EUCAST recommendations [87] using ETEST strips (BioMérieux). Briefly, overnight cultures of each strain to be tested were standardized to OD_600_ 0.063 in 0.85% NaCl (equivalent to McFarland standard 0.5) and distributed evenly across the surface of MH agar plates. E-test strips were placed on the surface of the plates, evenly spaced, and the plates were incubated for 18-24 hours at 37 °C. MICs were read according to the manufacturer’s instructions. MICs were also determined using the Broth Microdilution (BMD) method in accordance with the EUCAST recommendations [87] for specific β-lactams, as required, and for colistin sulphate (Acros Organics). Briefly, a series of antibiotic concentrations was prepared by two-fold serial dilution in MH broth in a clear-bottomed 96-well microtiter plate (Corning). The strain to be tested was added to the wells at approximately 5 x 10^5^ CFUs per well and plates were incubated for 18-24 hours at 37 °C. The MIC was defined as the lowest antibiotic concentration with no visible bacterial growth in the wells. When used for MIC assays, tazobactam was included at a fixed concentration of 4 μg/mL, in accordance with the EUCAST guidelines. All *S. maltophilia* MICs were performed in synthetic CF sputum medium (SCFM) as described in [88], using E-test strips (for β-lactam antibiotics) or the BMD method (for colistin). For *S. maltophilia* GUE, imipenem at a final concentration of 5 µg/mL was added to the overnight cultures to induce β-lactamase production.

The covalent DsbB inhibitor 4,5-dibromo-2-(2-chlorobenzyl)pyridazin-3(2H)-one [59] was used to chemically impair the function of the DSB system in *S. maltophilia* strains. Inactivation of DsbB results in abrogation of DsbA function [89] only in media free of small-molecule oxidants [90]. Therefore, MIC assays involving chemical inhibition of the DSB system were performed using SCFM media prepared as described in [88], except that L-cysteine was omitted. Either DMSO (vehicle control) or the covalent DsbB inhibitor 4,5-dibromo-2-(2-chlorobenzyl)pyridazin-3(2H)-one [59] (Bioduro-Sundia; ^1^H-NMR and LCMS spectra are provided in File S5), at a final concentration of 50 μM, were added to the cysteine-free SCFM medium, as required.

#### SDS-PAGE analysis and immunoblotting

Samples for immunoblotting were prepared as follows. Strains to be tested were grown on LB agar plates as lawns in the same manner as for MIC assays described above. Bacteria were collected using an inoculating loop and resuspended in LB to OD_600_ 2.0. The cell suspensions were centrifuged at 10,000 *x g* for 10 minutes and bacterial pellets were lysed by addition of BugBuster Master Mix (Merck Millipore) for 25 minutes at room temperature with gentle agitation. Subsequently, lysates were centrifuged at 10,000 *x g* for 10 minutes at 4 °C and the supernatant was added to 4 x Laemmli buffer. Samples were boiled for 5 minutes before separation by SDS-PAGE.

SDS-PAGE analysis was carried out using 10% BisTris NuPAGE gels (ThermoFisher Scientific) and MES/SDS running buffer prepared according to the manufacturer’s instructions; pre-stained protein markers (SeeBlue Plus 2, ThermoFisher Scientific) were included. Proteins were transferred to Amersham Protran nitrocellulose membranes (0.45 µm pore size, GE Life Sciences) using a Trans-Blot Turbo transfer system (Bio-Rad) before blocking in 3% w/v Bovine Serum Albumin (BSA)/TBS-T (0.1 % v/v Tween 20) or 5% w/v skimmed milk/TBS-T and addition of primary and secondary antibodies. The following primary antibodies were used in this study: Strep-Tactin-AP conjugate (Iba Lifesciences) (dilution 1:3,000 in 3 w/v % BSA/TBS-T), and mouse anti-DnaK 8E2/2 antibody (Enzo Life Sciences) (dilution 1:10,000 in 5% w/v skimmed milk/TBS-T). The following secondary antibodies were used in this study: goat anti-mouse IgG-AP conjugate (Sigma Aldrich) (dilution 1:6,000 in 5% w/v skimmed milk/TBS-T) and goat anti-mouse IgG-HRP conjugate (Sigma Aldrich) (dilution 1:6,000 in 5% w/v skimmed milk/TBS-T). Membranes were washed three times for 5 minutes with TBS-T prior to development. Development for AP conjugates was carried out using SigmaFast BCIP/NBT tablets.

Immunoblot samples were also analyzed for total protein content. SDS-PAGE analysis was carried out using 10% BisTris NuPAGE gels (ThermoFisher Scientific) and MES/SDS running buffer prepared according to the manufacturer’s instructions; pre-stained protein markers (SeeBlue Plus 2, ThermoFisher Scientific) were included. Gels were stained for total protein with SimplyBlue SafeStain (ThermoFisher Scientific) according to the manufacturer’s instructions.

#### β-Lactam hydrolysis assay

β-lactam hydrolysis measurements were carried out using the chromogenic β-lactam nitrocefin (Abcam). Briefly, overnight cultures of strains to be tested were centrifuged, pellets were weighed and resuspended in 150 μL of 100 mM sodium phosphate buffer (pH 7.0) per 1 mg of wet-cell pellet, and cells were lysed by sonication. Lysates were transferred into clear-bottomed 96-well microtiter plates (Corning) at volumes that corresponded to the following weights of bacterial cell pellets: strains harboring pDM1, pDM1-*bla*_L2-1_ and pDM1-*bla*_OXA-50_ (0.34 mg of cell pellet); strains harboring pDM1-*bla*_BEL-1_, pDM1-*bla*_AIM-1_ and pDM1-*bla*_SMB-1_ (0.17 mg of cell pellet); strains harboring pDM1-*bla*_POM-1_ (0.07 mg of cell pellet); strains harboring pDM1-*bla*_BPS-1m_ (0.07 mg of cell pellet); strains harboring pDM1-*bla*_CARB-2_ (0.03 mg of cell pellet). In all cases, nitrocefin was added at a final concentration of 400 μM and the final reaction volume was made up to 100 μL using 100 mM sodium phosphate buffer (pH 7.0). Nitrocefin hydrolysis was monitored at 25 °C by recording absorbance at 490 nm at 60-second intervals for 15 minutes using an Infinite M200 Pro microplate reader (Tecan). The amount of nitrocefin hydrolyzed by each lysate in 15 minutes was calculated using a standard curve generated by acid hydrolysis of nitrocefin standards.

#### Galleria mellonella survival assay

The wax moth model *G. mellonella* was used for *in vivo* survival assays [91]. Individual *G. mellonella* larvae were randomly allocated to experimental groups; no masking was used. Overnight cultures of all the strains to be tested were standardized to OD_600_ 1.0, suspensions were centrifuged, and the pellets were washed three times in PBS and serially diluted. For experiments with *P. aeruginosa* G6R7, 10 μL of the 1:10,000 dilution of each bacterial suspension was injected into the last right abdominal proleg of 40 *G. mellonella* larvae per condition. One hour after infection, larvae were injected with 2.75 μL of piperacillin to a final concentration of 5 μg/mL in the last left abdominal proleg. For experiments with *P. aeruginosa* CDC #773 10 μL of the 1:1,000 dilution of each bacterial suspension was injected into the last right abdominal proleg of 30 *G. mellonella* larvae per condition. Immediately after the injection with the inoculum, the larvae were injected with 4.5 μl of ceftazidime to a final concentration of 6.5 μg/mL in the last left abdominal proleg. All larvae were incubated at 37 °C and their mortality was monitored for 30 hours. Death was recorded when larvae turned black due to melanization and did not respond to physical stimulation. For each experiment, an additional ten larvae were injected with PBS as negative control and experiments were discontinued and discounted if mortality was greater than 10% in the PBS control.

#### S. maltophilia – P. aeruginosa protection assay

The protection assay was based on the approach described in [15]. Briefly, 75 μL of double-strength SCFM medium were transferred into clear-bottomed 96-well microtiter plates (VWR) and inoculated with *S. maltophilia* AMM or its *dsbA dsbL* mutant that had been grown in SCFM medium at 37 °C overnight; *S. maltophilia* strains were inoculated at approximately 5 x 10^4^, as appropriate. Plates were incubated at 37 °C for 6 hours. Double-strength solutions of ceftazidime at decreasing concentrations were prepared by two-fold serial dilution in sterile ultra-pure H_2_O, and were added to the wells, as required. *P. aeruginosa* PA14 was immediately added to all the wells at approximately 5 x 10^4^ CFUs, and the plates were incubated for 20 hours at 37 °C.

To enumerate *P. aeruginosa* in this experiment, the *P. aeruginosa* PA14 *att*Tn*7::accC* strain was used. Following the 20-hour incubation step, serial dilutions of the content of each well were performed in MH broth down to a 10^-7^ dilution, plated on MH agar supplemented with gentamicin (*S. maltophilia* AMM strains are sensitive to gentamicin, whereas *P. aeruginosa* PA14 *att*Tn*7::accC* harbours a gentamicin resistance gene on its Tn*7* site) and incubated at 37 °C overnight. CFUs were enumerated the following day. To enumerate *S. maltophilia* in this experiment*, S. maltophilia* AMM *att*Tn*7::accC msfgfp* or its *dsbA dsbL* mutant were used. Following the 20-hour incubation step, serial dilutions of the content of each well were performed in MH broth down to a 10^-7^ dilution, plated on MH agar supplemented with gentamicin (*S. maltophilia* AMM strains harbour a gentamicin resistance gene on their Tn*7* site, whereas *P. aeruginosa* PA14 is sensitive to gentamicin) and incubated at 37 °C overnight. CFUs were enumerated the following day.

#### Statistical analysis of experimental data

The total number of performed biological experiments and technical repeats are mentioned in the figure legend of each display item. Biological replication refers to completely independent repetition of an experiment using different biological and chemical materials. Technical replication refers to independent data recordings using the same biological sample.

Antibiotic MIC values were determined in biological triplicate, except for MIC values recorded for *dsbA* complementation experiments in our *E. coli* K-12 inducible system that were carried out in duplicate. All ETEST MICs were determined as a single technical replicate, and all BMD MICs were determined in technical triplicate. All recorded MIC values are displayed in the relevant graphs; for MIC assays where three or more biological experiments were performed, the bars indicate the median value, while for assays where two biological experiments were performed the bars indicate the most conservative of the two values (i.e., for increasing trends, the value representing the smallest increase and for decreasing trends, the value representing the smallest decrease). We note that in line with recommended practice, our MIC results were not averaged. This should be avoided because of the quantized nature of MIC assays, which only inform on bacterial survival for specific antibiotic concentrations and do not provide information for antibiotic concentrations that lie in-between the tested values.

For all other assays, statistical analysis was performed in GraphPad Prism v8.3.1 using either an unpaired T-test with Welch’s correction, or a Mantel-Cox logrank test, as appropriate. Statistical significance was defined as p < 0.05. Outliers were defined as any technical repeat >2 SD away from the average of the other technical repeats within the same biological experiment. Such data were excluded and all remaining data were included in the analysis. Detailed information for each figure is provided below:

Figure 2B: unpaired T-test with Welch’s correction; n=3; 3.417 degrees of freedom, t-value=0.3927, p=0.7178 (non-significance) (for pDM1 strains); 2.933 degrees of freedom, t-value=0.3296, p=0.7639 (non-significance) (for pDM1-*bla*_L2-1_ strains); 2.021 degrees of freedom, t-value=7.549, p=0.0166 (significance) (for pDM1-*bla*_BEL-1_ strains); 2.146 degrees of freedom, t-value=9.153, p=0.0093 (significance) (for pDM1-*bla*_CARB-1_ strains); 2.320 degrees of freedom, t-value=5.668, p=0.0210 (significance) (for pDM1-*bla*_AIM-1_ strains); 3.316 degrees of freedom, t-value=4.353, p=0.0182 (significance) (for pDM1-*bla*_OXA-50_ strains); 3.416 degrees of freedom, t-value=13.68, p=0.0004 (significance) (for pDM1-*bla*_BPS-1m_ strains).

Figure 3C: Mantel-Cox test; n=40; p=0.3173 (non-significance) (*P. aeruginosa* vs *P. aeruginosa dsbA1*), p<0.0001 (significance) (*P. aeruginosa* vs *P. aeruginosa* treated with piperacillin), p<0.0001 (significance) (*P. aeruginosa dsbA1* vs *P. aeruginosa* treated with piperacillin), p=0.0147 (significance) (*P. aeruginosa* treated with piperacillin vs *P. aeruginosa dsbA1* treated with piperacillin).

Figure 3F: Mantel-Cox test; n=30; p<0.0001 (significance) (*P. aeruginosa* vs *P. aeruginosa dsbA1*), p>0.9999 (non-significance) (*P. aeruginosa* vs *P. aeruginosa* treated with ceftazidime), p<0.0001 (significance) (*P. aeruginosa dsbA1* vs *P. aeruginosa* treated with ceftazidime), p<0.0001 (significance) (*P. aeruginosa* treated with ceftazidime vs *P. aeruginosa dsbA1* treated with ceftazidime).

Figure S4C: unpaired T-test with Welch’s correction; n=3; 3.417 degrees of freedom, t-value=0.3927, p=0.7178 (non-significance) (for pDM1 strains); 2.933 degrees of freedom, t-value=0.3296, p=0.7639 (non-significance) (for pDM1-*bla*_L2-1_ strains); 3.998 degrees of freedom, t-value=4.100, p=0.0149 (significance) (for pDM1-*bla*_POM-1_ strains); 2.345 degrees of freedom, t-value=15.02, p=0.0022 (significance) (for pDM1-*bla*_SMB-1_ strains).

#### Bioinformatics

The following bioinformatics analyses were performed in this study. Short scripts and pipelines were written in Perl (version 5.18.2) and executed on macOS Sierra 10.12.5.

#### β-lactamase enzymes

All available protein sequences of β-lactamases were downloaded from http://www.bldb.eu [81] (29 November 2024). Sequences were clustered using the ucluster software with a 90% identity threshold and the cluster_fast option (USEARCH v.7.0 [92]); the centroid of each cluster was used as a cluster identifier for every sequence. All sequences were searched for the presence of cysteine residues using a Perl script. Proteins with two or more cysteines after the first 30 amino acids of their primary sequence were considered potential substrates of the DSB system for organisms where oxidative protein folding is carried out by DsbA and provided that translocation of the β-lactamase outside the cytoplasm is performed by the Sec system. The first 30 amino acids of each sequence were excluded to avoid considering cysteines that are part of the signal sequence mediating the translocation of these enzymes outside the cytoplasm. The results of the analysis can be found in File S1.

#### Stenotrophomonas MCR-like enzymes

Hidden Markov Models built with validated sequences of MCR-like and EptA-like proteins were used to identify MCR analogues in a total of 106 complete genomes of the *Stenotrophomonas* genus, downloaded from the NCBI repository (30 March 2023). The analysis was performed with *hmmsearch* (HMMER v.3.1b2) [93] and only hits with evalues < 1e-10 were considered. The 146 obtained sequences were aligned using MUSCLE [94] and a phylogenetic tree was built from the alignment using FastTree 2.1.7 with the wag substitution matrix and the gamma option [95]. The assignment of each MCR-like protein sequence to a specific phylogenetic group was carried out based on the best fitting *hmmscan* model. The results of the analysis can be found in File S4.

#### Data availability

All data generated during this study that support the findings are included in the manuscript or the Supplementary Information. All materials are available from the corresponding author upon request.

## ACKNOWLEDGEMENTS

We thank L. Dortet for the kind gift of any *P. aeruginosa* and *S. maltophilia* clinical isolates that do not originate from the Centers for Disease Control and Prevention. Publication of this work was supported by the National Institute of Allergy and Infectious Diseases of the National Institutes of Health under Award Number R01AI158753 (to D.A.I.M.); the content is solely the responsibility of the authors and does not necessarily represent the official views of the National Institutes of Health. Additionally, this study was funded by the Medical Research Council Career Development Award MR/M009505/1 (to D.A.I.M.), a Texas Biologics (TXBio) grant (Award Number TXB-24-02) from The Cockrell School of Engineering at The University of Texas at Austin (to D.A.I.M.), the UT | Portugal Extra Exploratory Project grant 2022.15740.UTA from the Fundação para a Ciência e a Tecnologia, I.P. (to D.A.I.M.), and the Welch Foundation grant F-2250-20250403 (to D.A.I.M.), the institutional Biotechnology and Biological Sciences Research Council (BBSRC)-Doctoral Training Program studentship BB/M011178/1 (to N.K.), the MCIN/AEI/10.13039/501100011033 Spanish agency through the Ramon y Cajal RYC2019-026551-I and PID2021-123000OB-I00 grants (to P.B.), the Indiana University Bloomington start-up funds (to C.L.), and the Cystic Fibrosis Foundation through the Pilot and Feasibility Award 004846I222 (to C.L.), the Swiss National Science Foundation Ambizione Fellowship PZ00P3_180142 (to D.G.), as well as the NC3Rs grant NC/V001582/1 (to E.M. and R.R.MC.), the BBSRC New Investigator Award BB/V007823/1 (to R.R.MC.), the Academy of Medical Sciences / the Wellcome Trust / the Government Department of Business, Energy and Industrial Strategy / the British Heart Foundation / Diabetes UK Springboard Award SBF006\1040 (to R.R.MC.), and the Medical Research Council grant MR/Y001354/1 (to R.R.MC.).

## AUTHOR CONTRIBUTIONS

N.K., R.C.D.F. and D.A.I.M. designed the research. N.K. performed most of the experiments. P.B. and A.F. provided strains, genetic tools and advice on *P. aeruginosa* molecular biology. K.E.P. designed and constructed plasmids used to complement *P. aeruginosa* and *S. maltophilia* clinical strains. C.L. provided materials and advice on the chemical inhibition of the DSB system. D.G. performed *in silico* analyses and advised on several aspects of the project. L.E., E.M and R.R.MC performed *G. mellonella* survival assays. N.K., R.C.D.F. and D.A.I.M. wrote the manuscript with input from all authors. D.A.I.M. directed the project.

## SUPPLEMENTARY INFORMATION FOR

### This PDF file includes

Figures S1 to S5

Tables S1 to S5

Legends for Files S1 to S5

Supplementary references

## SUPPLEMENTARY FIGURES

**Figure S1.**
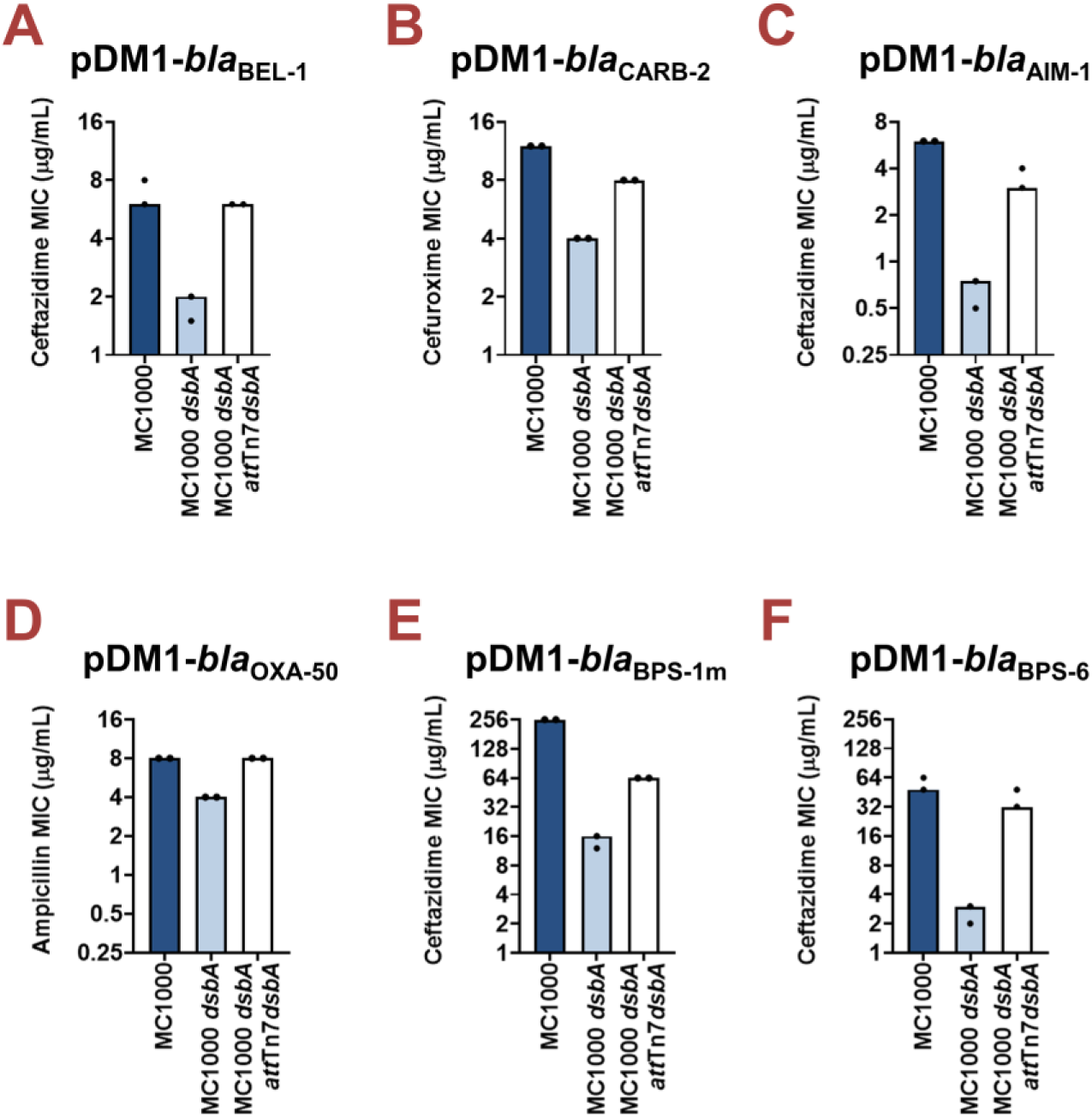
Complementation of *dsbA* restores the β-lactam MIC values for *E. coli* MC1000 *dsbA* expressing β-lactamase enzymes. Re-insertion of *dsbA* at the *att*Tn*7* site of the chromosome restores representative β-lactam MIC values for *E. coli* MC1000 *dsbA* harboring **(A)** pDM1-*bla*_BEL-1_ (ceftazidime MIC), **(B)** pDM1-*bla*_CARB-2_ (cefuroxime MIC), **(C)** pDM1-*bla*_AIM-1_ (ceftazidime MIC), **(D)** pDM1-*bla*_OXA-50_ (ampicillin MIC), **(E)** pDM1-*bla*_BPS-1m_ (ceftazidime MIC), and **(F)** pDM1-*bla*_BPS-6_ (ceftazidime MIC). Graphs show MIC values (µg/mL) and are representative of two biological experiments, each conducted as a single technical repeat.

**Figure S2.**
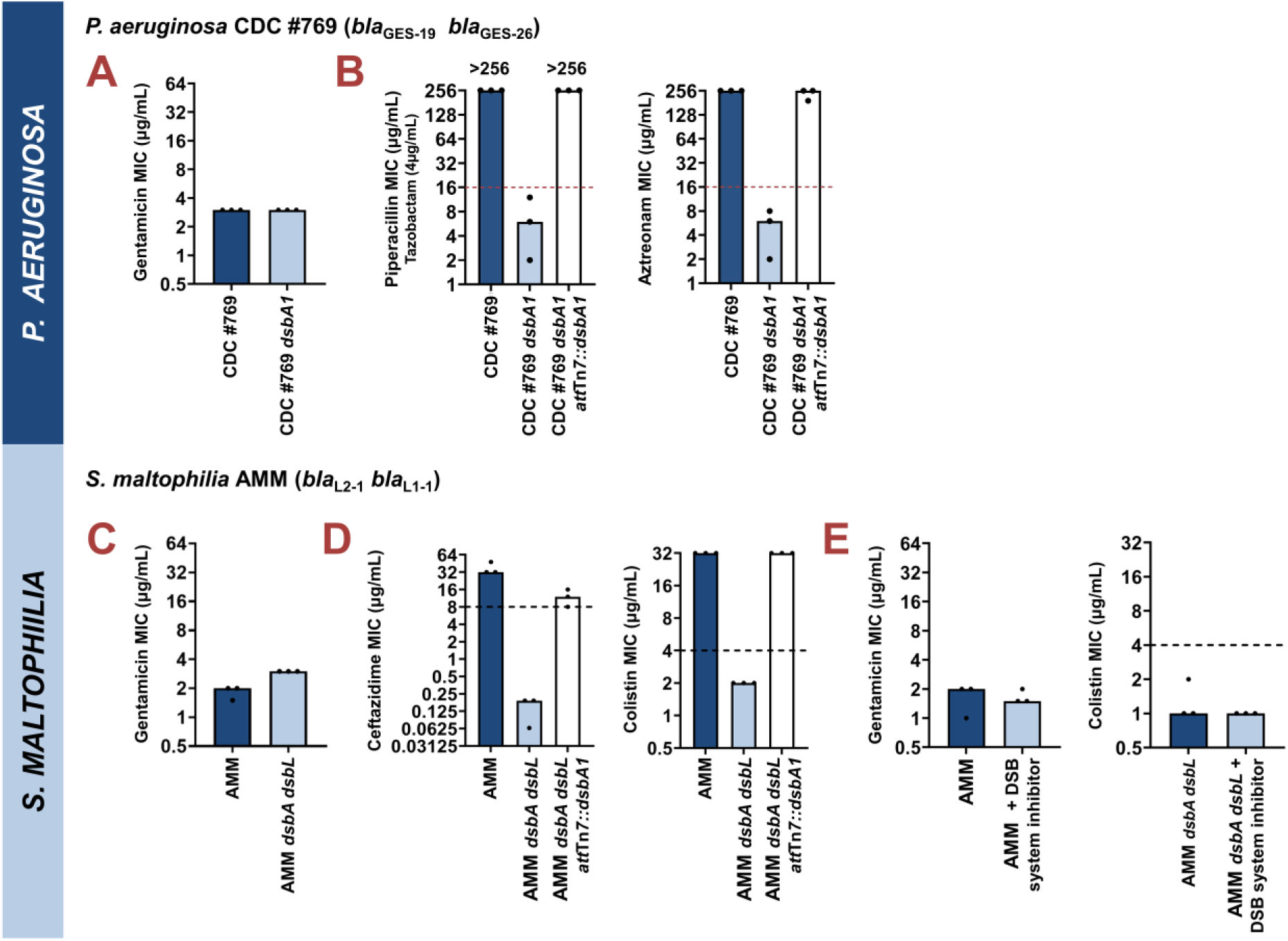
Assessment of off-target effects for clinical strains of *P. aeruginosa* and *S. maltophilia* that are deficient in oxidative protein folding. (A) *P. aeruginosa* CDC #769 and its mutant lacking *dsbA1* have identical gentamicin MIC values, confirming that absence of DsbA does not compromise the general ability of the strain to resist antibiotic stress. **(B)** Re-insertion of the *dsbA1* gene from *P. aeruginosa* PAO1 at the *att*Tn*7* site of the chromosome restores representative antibiotic MIC values for *P. aeruginosa* CDC #769 *dsbA1* (left, piperacillin/tazobactam MIC; right, aztreonam MIC). (C) *S. maltophilia* AMM and its mutant lacking *dsbA* and *dsbL* have near-identical gentamicin MIC values, confirming that absence of DsbA and DsbL does not compromise the general ability of the strain to resist antibiotic stress. **(D)** Re-insertion of the *dsbA1* gene from *P. aeruginosa* PAO1 at the *att*Tn*7* site of the chromosome restores representative antibiotic MIC values for *S. maltophilia* AMM *dsbA dsbL* (left, ceftazidime MIC; right, colistin MIC). **(E)** Changes in MIC values observed using the DSB system inhibitor (compound 36) are due solely to inhibition of the DSB system. The gentamicin MIC value of *S. maltophilia* AMM remains unchanged upon addition of the inhibitor (left), and the same is observed for the colistin MIC value of *S. maltophilia* AMM *dsbA dsbL* in the presence of the compound (right). This indicates that the chemical compound used in this study only affects the function of the DSB system proteins. For all panels, graphs show MIC values (µg/mL) and are representative of three biological experiments. β-Lactam MICs were conducted as a single technical repeat and colistin MICs were conducted in technical triplicate; red dotted lines indicate the EUCAST clinical breakpoint for each antibiotic, where applicable. In the absence of EUCAST clinical breakpoints for *S. maltophilia*, the black dotted lines indicate the EUCAST clinical breakpoint for each antibiotic for the related pathogen *P. aeruginosa*, where applicable.

**Figure S3.**
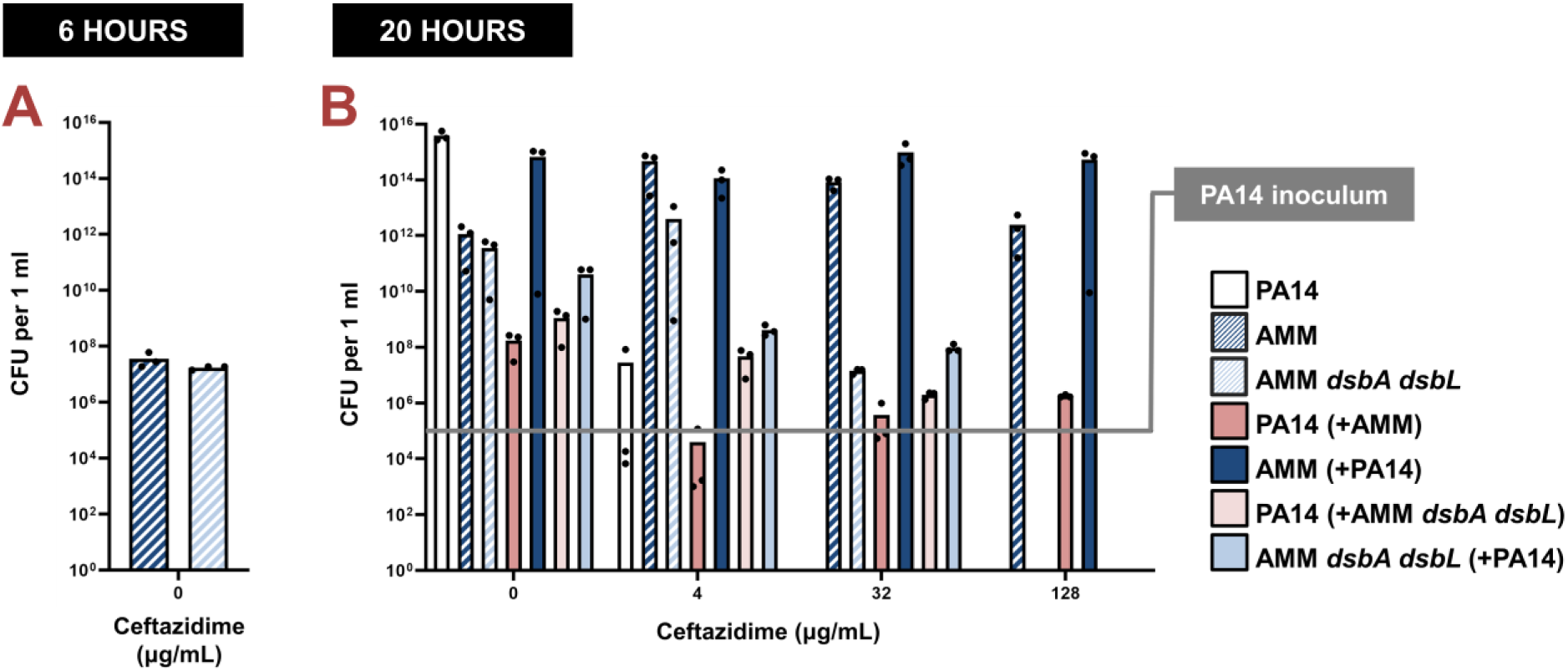
Protection of *P. aeruginosa* by *S. maltophilia* clinical strains is dependent on oxidative protein folding. **(A)** Comparison of the colony forming units (CFUs) of *S. maltophilia* AMM with the CFUs of *S. maltophilia* AMM *dsbA dsbL* after six hours of growth, prior to *P. aeruginosa* PA14 addition. The two *S. maltophilia* strains display equivalent growth. **(B)** Complementary analysis to Fig. 4E; here the CFUs of all *P. aeruginosa* and *S. maltophilia* strains were enumerated in isolation and in mixed culture conditions for a more limited set of antibiotic concentrations. Equivalent trends to Fig. 4E are observed. The susceptible *P. aeruginosa* strain PA14 can survive exposure to ceftazidime up to a maximum concentration of 4 μg/mL when cultured in isolation (white bars). By contrast, if co-cultured in the presence of *S. maltophilia* AMM (dark blue bars), which can hydrolyze ceftazidime through the action of its L1-1 β-lactamase enzyme, *P. aeruginosa* PA14 (dark pink bars) can survive and actively grow in higher concentrations of ceftazidime (see 128 μg/mL of ceftazidime). This protection is abolished if *P. aeruginosa* PA14 (light pink bars) is co-cultured with *S. maltophilia* AMM *dsbA dsbL* (light blue bars). In this case, L1-1 is inactive (as shown in Fig. 4AB and [1]), resulting in killing of *S. maltophilia* AMM and, in turn, eradication of *P. aeruginosa* PA14 (see 128 μg/mL of ceftazidime, absence of light pink bars). Three biological replicates were conducted in technical triplicate and mean CFU values are shown. The grey line indicates the *P. aeruginosa* PA14 inoculum. The mean CFU values used to generate this figure are presented in File S2C.

**Figure S4.**
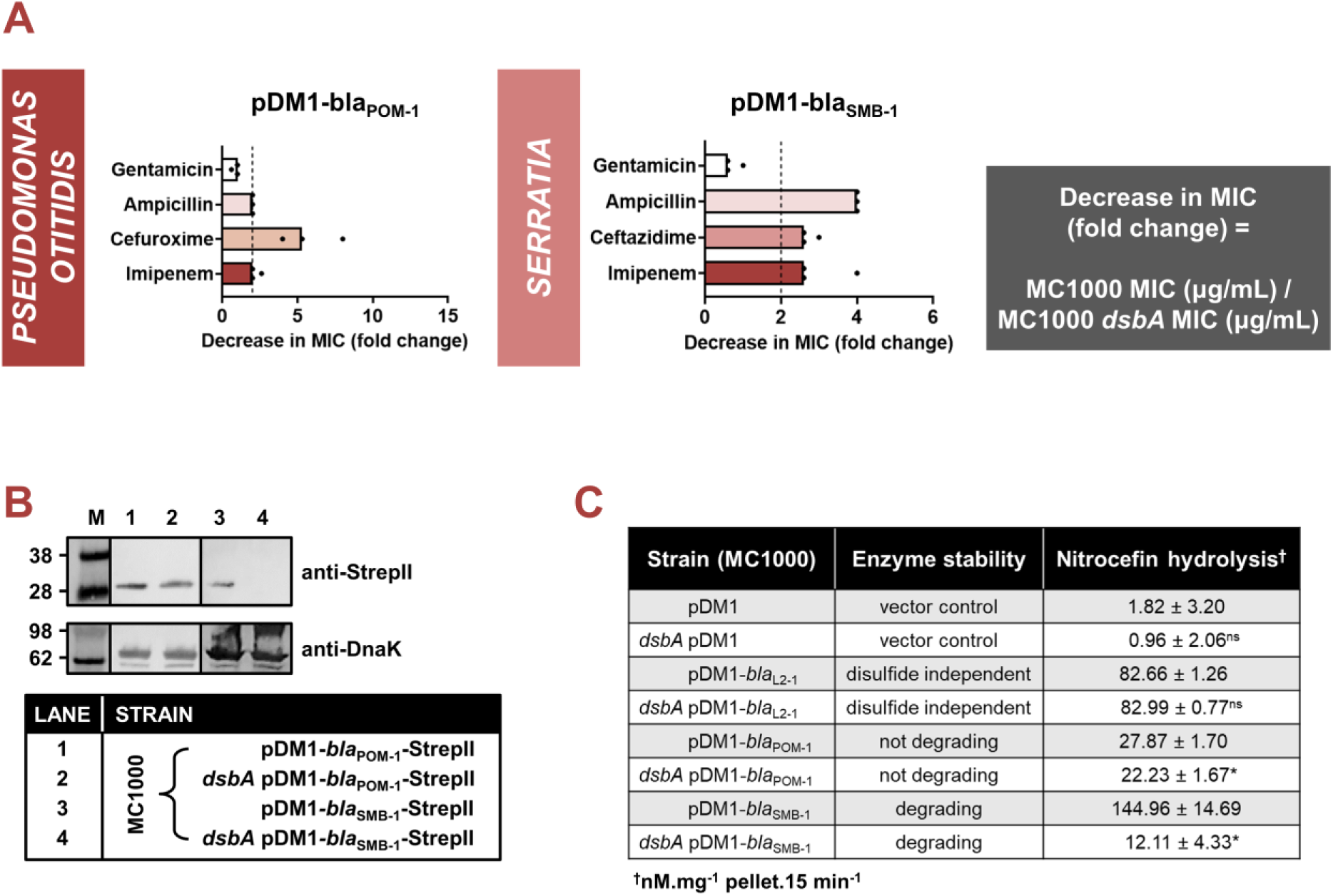
The activity of additional species-specific β-lactamases depends on disulfide bond formation. **(A)** β-Lactam MIC values for *E. coli* MC1000 expressing disulfide-bond-containing β-lactamases from *P. otitidis* (left, POM-1; Table S1) and *Serratia spp*. (right, SMB-1; Table S1) are reduced in the absence of DsbA (MIC fold changes: >2; fold change of 2 is indicated by the black dotted lines). No changes in MIC values are observed for the aminoglycoside antibiotic gentamicin (white bars) confirming that absence of DsbA does not compromise the general ability of this strain to resist antibiotic stress. Graphs show MIC fold changes for β-lactamase-expressing *E. coli* MC1000 and its *dsbA* mutant. MIC assays were performed in three biological experiments each conducted as a single technical repeat; the MIC values used to generate this figure are presented in File S2A (rows 22-25). **(B)** Protein levels of disulfide-bond-containing β-lactamases are either unaffected (POM-1) or drastically reduced (SMB-1) when these enzymes are expressed in *E. coli* MC1000 *dsbA*. Protein levels of StrepII-tagged β-lactamases were assessed using a Strep-Tactin-AP conjugate. A representative blot from three biological experiments, each conducted as a single technical repeat, is shown; molecular weight markers (M) are on the left, DnaK was used as a loading control and solid black lines indicate where the membrane was cut. Full immunoblots and SDS PAGE analysis of the immunoblot samples for total protein content are shown in File S3. **(C)** The hydrolytic activities of both tested β-lactamases are significantly reduced in the absence of DsbA. The hydrolytic activities of strains harboring the empty vector or expressing the control enzyme L2-1 show no dependence on DsbA; the same data for the control strains are also shown in Fig. 2B. n=3 (each conducted in technical duplicate), table shows means ± SD, significance is indicated by * = p < 0.05, ns = non-significant.

**Figure S5.**
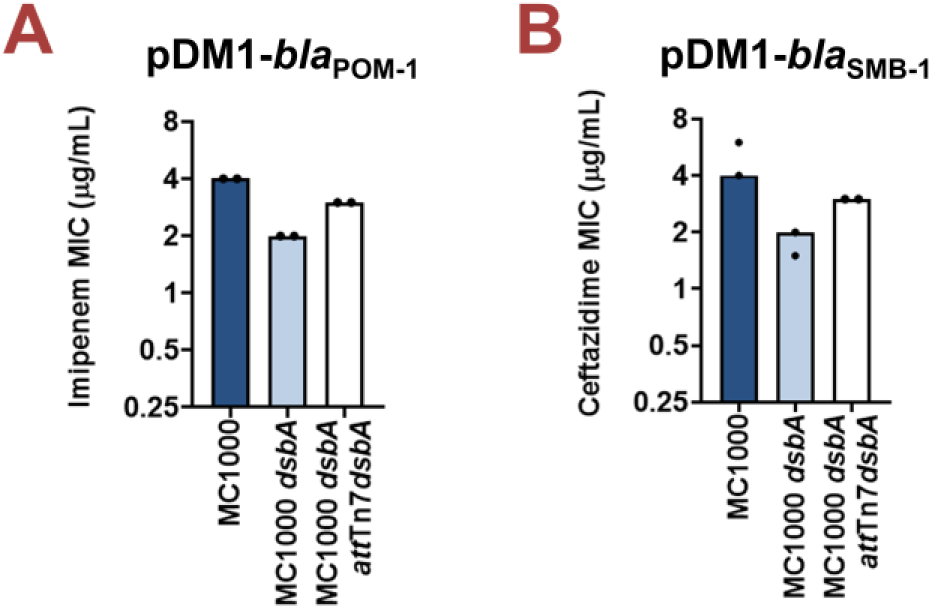
Complementation of *dsbA* restores the β-lactam MIC values for *E. coli* MC1000 *dsbA* expressing β-lactamases. Re-insertion of *dsbA* at the *att*Tn*7* site of the chromosome restores representative β-lactam MIC values for *E. coli* MC1000 *dsbA* harboring **(A)** pDM1-*bla*_POM-1_ (imipenem MIC), and **(B)** pDM1-*bla*_SMB-1_ (ceftazidime MIC). Graphs show MIC values (µg/mL) and are representative of two biological experiments, each conducted as a single technical repeat.

## SUPPLEMENTARY TABLES

**Table S1.** Overview of the β-lactamase enzymes investigated in this study. All tested enzymes belong to distinct phylogenetic clusters (see File S1), with the exception of BPS-1m and BPS-6. The “Cysteine positions” column states the positions of cysteine residues after amino acid 30 and hence, does not include amino acids that are part of the periplasmic signal sequence which is cleaved after protein translocation. All β-lactamase enzymes except L2-1 and LUT-1 (shaded in grey), which are used as negative controls throughout this study, have one or more disulfide bonds. Both L2-1 and LUT-1 contain two or more cysteine residues, but lack disulfide bonds as they are transported to the periplasm in a folded state by the Twin-arginine translocation (Tat) system; for L2-1 Tat-dependent translocation has been experimentally confirmed [2], whereas for LUT-1 this is strongly corroborated by signal peptide prediction software (SignalP 5.0 [3] likelihood scores: Sec/SPI = 0.0572, Tat/SPI = 0.9312, Sec/SPII (lipoprotein) = 0.0087, other = 0.0029). The “Mob.” (mobilizable) column refers to the possibility for the β-lactamase gene to be mobilized from the chromosome; “yes” indicates that the gene of interest is located on a mobile element, while “no” refers to immobile chromosomally-encoded enzymes. The “Spectrum” column refers to the hydrolytic spectrum of each tested enzyme; tested enzymes are narrow-spectrum β-lactamases (NS), extended-spectrum β-lactamases (ESBL) or carbapenemases. The “Inh.” (inhibition) column refers to classical inhibitor susceptibility i.e., susceptibility to inhibition by clavulanic acid, tazobactam or sulbactam. Finally, the “Organism” column refers to the bacterial species that most commonly express the tested β-lactamase enzymes.

| ENZYME | CYSTEINE POSITIONS | AMBLER CLASS | MOB. | SPECTRUM | INH. | ORGANISM |
| --- | --- | --- | --- | --- | --- | --- |
| **L2-1** | C82 C136 C233 | A | no | ESBL | yes | *Stenotrophomonas maltophilia* |
| **LUT-1** | C54 C129 | A | no [4] | NS | yes | *Pseudomonas luteola* |
| **BEL-1** | C61 C231 | A | yes [5] | ESBL | yes | *Pseudomonas aeruginosa* |
| **CARB-2** | C72 C118 | A | yes [6] | NS | yes | *Pseudomonas spp.* |
| **BPS-1m** | C75 C83 C129 | A | no [7] | ESBL | yes | *Burkholderia pseudomallei* |
| **BPS-6** | C75 C83 C129 | A | no [8] | ESBL | yes | *Burkholderia pseudomallei* |
| **GES-19, 20, or 26** | C63 C233 | A | yes [9] | ESBLs | yes | *Enterobacteriaceae, Pseudomonas aeruginosa* |
| **AIM-1** | C31 C56 C194 C199 C234 C274 | B3 | yes [10] | carbapenemase | no [11] | *Pseudomonas aeruginosa* |
| **L1-1** | C239 C265 | B3 | no [11] | carbapenemase | no [11] | *Stenotrophomonas maltophilia* |
| **POM-1** | C237 C265 | B3 | no [12] | carbapenemase | no [11] | *Pseudomonas otitidis* |
| **SMB-1** | C180 C 185 C226 C260 | B3 | yes [13] | carbapenemase | no [11] | *Serratia spp.* |
| **OXA-50** | C208 C211 | D | no [14] | NS | no [14] | *Pseudomonas spp.* |

**Table S2.**
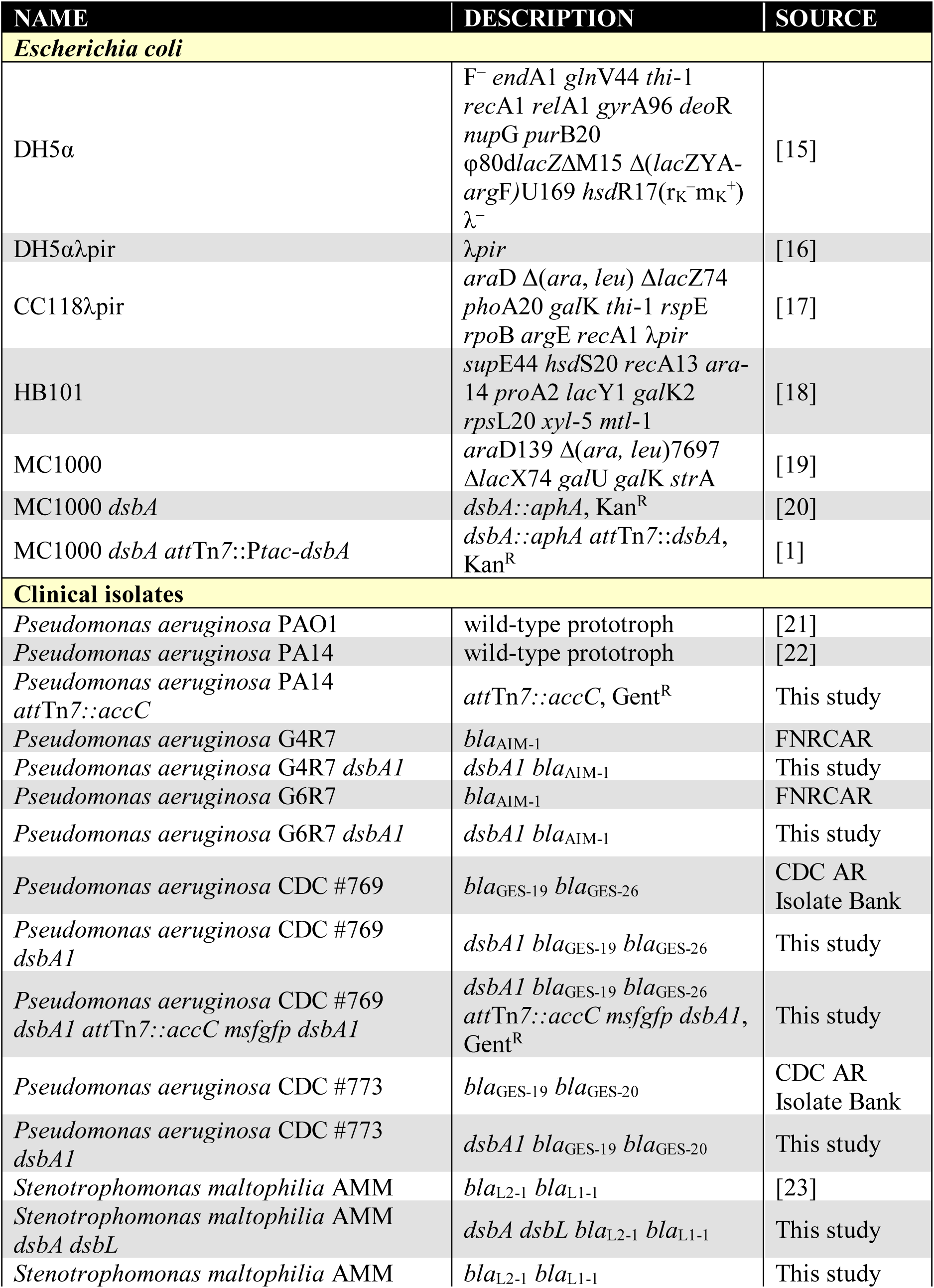
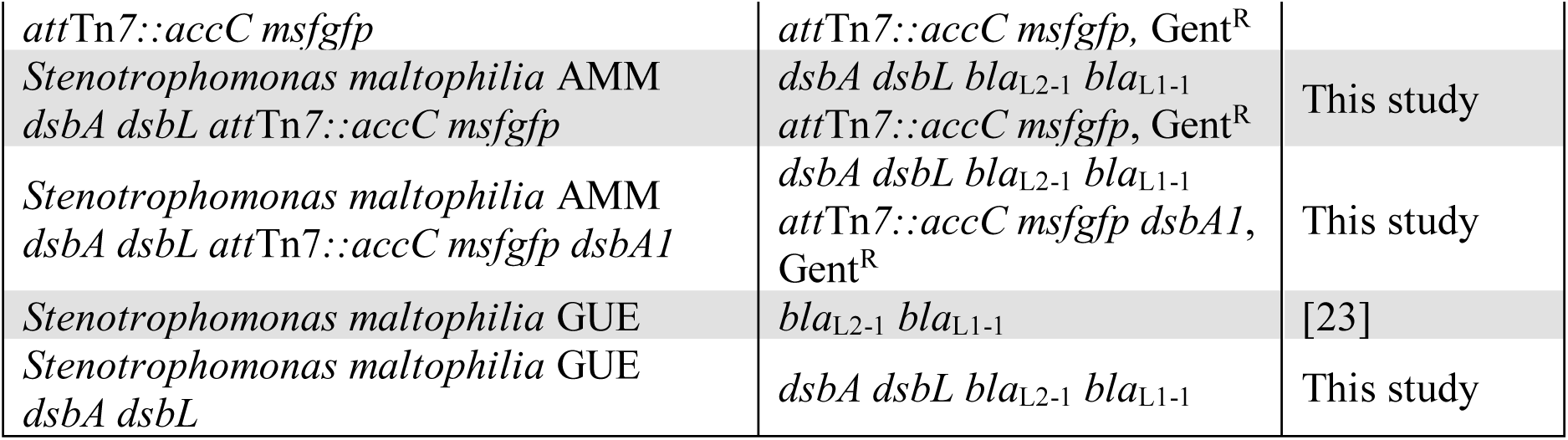
Bacterial strains used in this study. All listed isolates are clinical strains. “FNRCAR” refers to the French National Reference Centre for Antibiotic Resistance in Le Kremlin-Bicêtre, France, and “CDC AR Isolate bank” refers to the Centers for Disease Control and Prevention Antibiotic Resistance Isolate Bank in Atlanta, GA, USA.

**Table S3.**
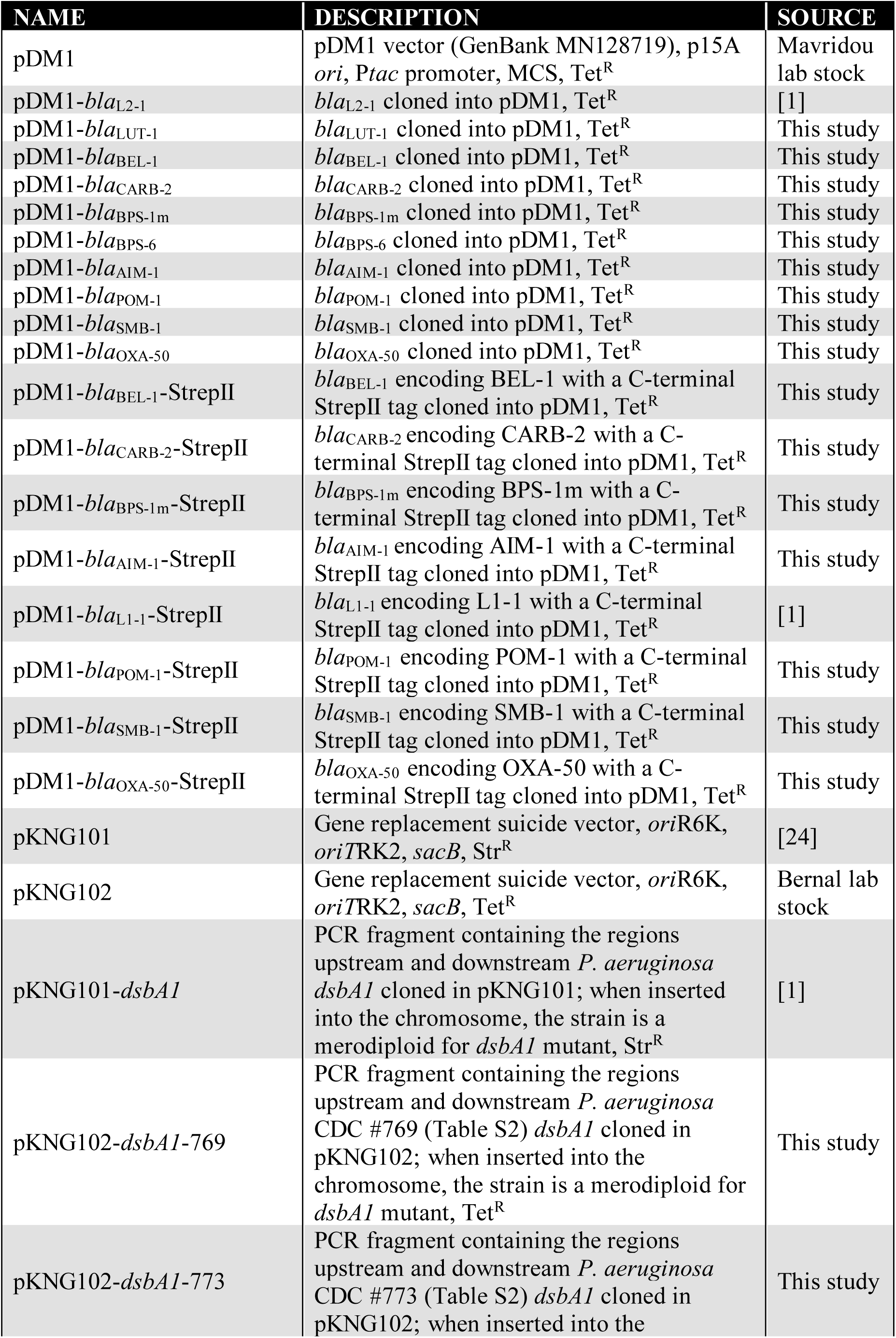
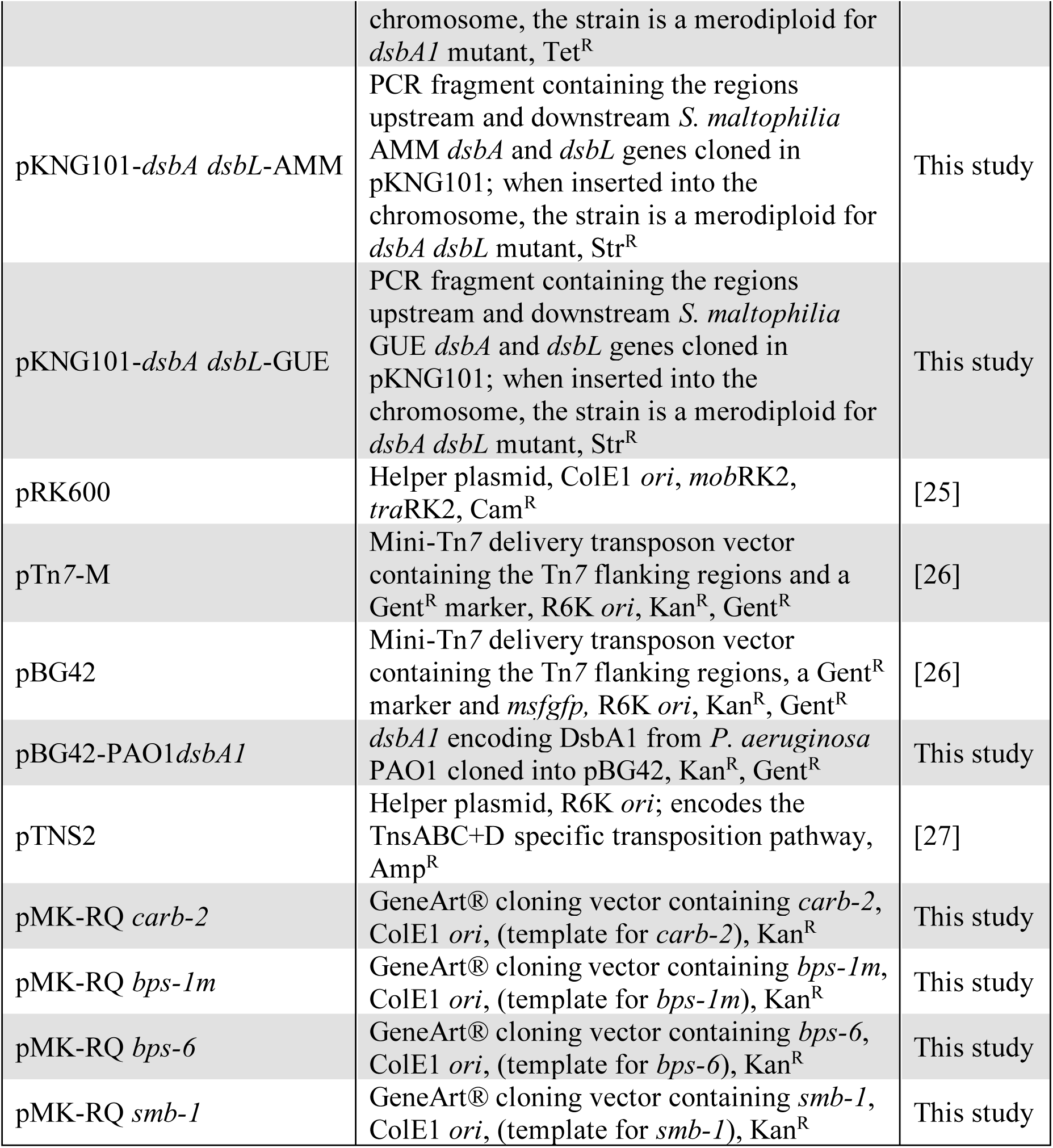
Plasmids used in this study.

**Table S4.**
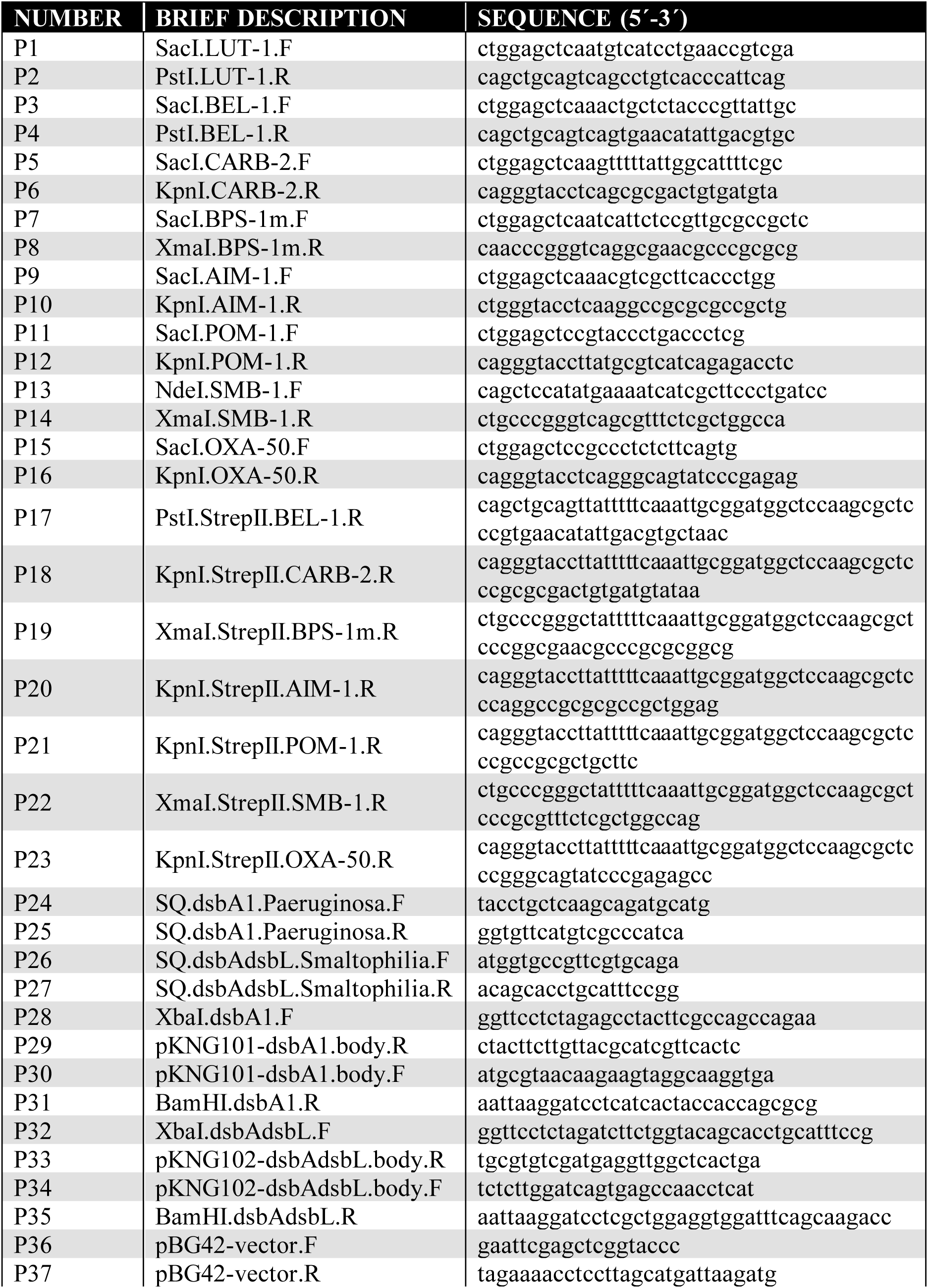
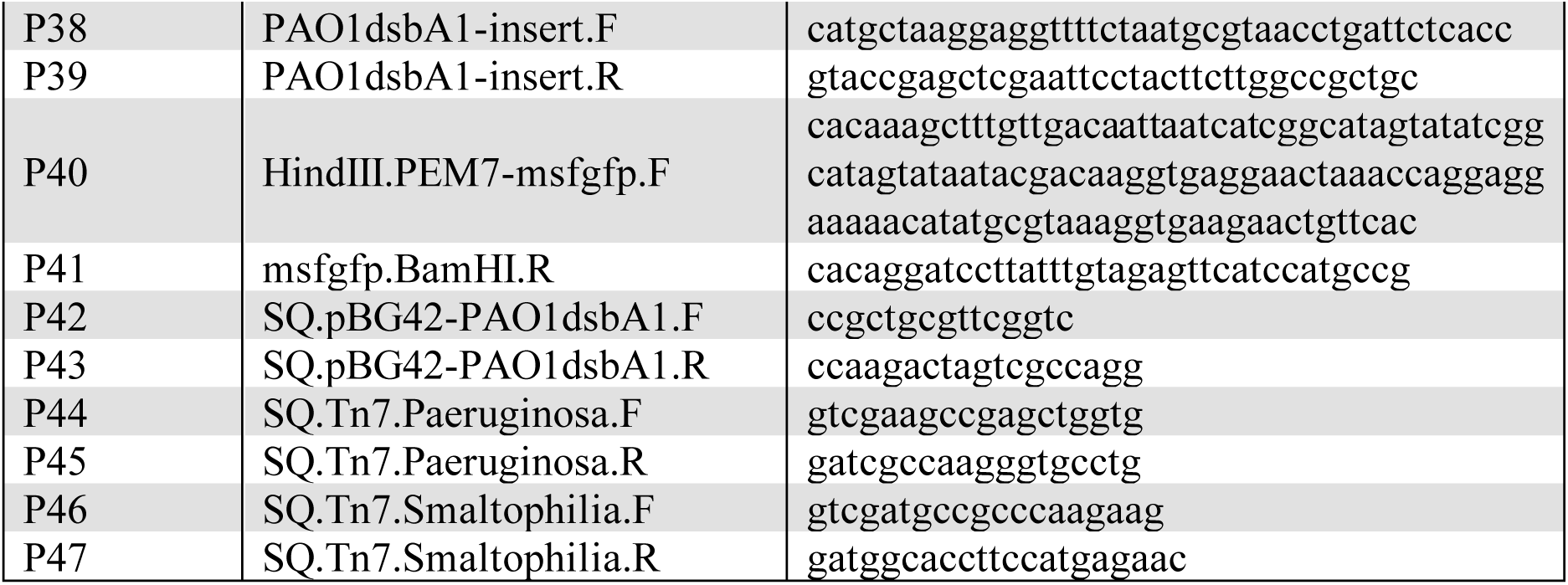
Oligonucleotide primers used in this study. The “Brief description” column provides basic information on the primer design (restriction enzyme used for cloning, encoded protein or gene replaced by antibiotic resistance cassette, forward or reverse orientation of the primer (F or R); SQ stands for sequencing primers).

**Table S5.** Sources of genomic DNA used for amplification of β-lactamase genes used in this study. CRBIP stands for Centre de Ressources Biologiques de l’Institut Pasteur, France and FNRCAR refers to the French National Reference Centre for Antibiotic Resistance in Le Kremlin-Bicêtre, France.

| STRAIN | GENE | SOURCE |
| --- | --- | --- |
| *Pseudomonas aeruginosa* 51170 | *bla*<sub>BEL-1</sub> | [5] |
| *Pseudomonas luteola* CIP 102067 | *bla*<sub>LUT-1</sub> | CRBIP |
| *Pseudomonas aeruginosa* G4R7 | *bla*<sub>AIM-1</sub> | FNRCAR |
| *Pseudomonas otitidis* CIP 109236T | *bla*<sub>POM-1</sub> | CRBIP |
| *Pseudomonas aeruginosa* PAO1 LA | *bla*<sub>OXA-50</sub> | [21] |

## LEGENDS FOR SUPPLEMENTARY DATA FILES

**File S1. Analysis of the cysteine content and phylogeny of all identified β-lactamases.** 7,741 unique β-lactamase protein sequences were clustered with a 90% identity threshold and the centroid of each cluster was used as a phylogenetic cluster identifier for each sequence (“Phylogenetic cluster (90% ID)” column). All sequences were searched for the presence of cysteine residues (“Total number of cysteines” and “Positions of all cysteines” columns). Proteins with two or more cysteines after the first 30 amino acids of their primary sequence (cells shaded in grey in the “Number of cysteines after position 30” column) are potential substrates of the DSB system for organisms where oxidative protein folding is carried out by DsbA and provided that translocation of the β-lactamase outside the cytoplasm is performed by the Sec system. The first 30 amino acids of each sequence were excluded to avoid considering cysteines that are part of the signal sequence mediating the translocation of these enzymes outside the cytoplasm. Cells shaded in grey in the “Reported in pathogens” column mark β-lactamases that are found in pathogens or organisms capable of causing opportunistic infections. The Ambler class of each enzyme is indicated in the “Ambler class column” and each class (A, B1, B2, B3, C and D) is highlighted in a different color.

**File S2. Data used to generate Fig. 1, Fig. S4, Fig. 4B and Fig. S3B. (A)** MIC values (µg/mL) used to generate Fig. 1 are in rows 2-7 [strains serving as negative controls; *E. coli* MC1000 strains harboring pDM1 (vector alone), pDM1-*bla*_L2-1_ or pDM1-*bla*_LUT-1_ (cysteine-containing β-lactamases which lack disulfide bonds)] and rows 9-20. MIC values (µg/mL) used to generate Fig. S4 are in rows 22-25. The aminoglycoside antibiotic gentamicin serves as a negative control for all strains. Cells marked with a dash (-) represent strain-antibiotic combinations that were not tested. **(B)** *P. aeruginosa* PA14 colony forming unit (CFU) counts used to generate Fig. 4E. **(C)** *P. aeruginosa* PA14, *S. maltophilia* AMM and *S. maltophilia AMM dsbA dsbL* CFU counts used to generate Fig. S3B. For all tabs, three biological experiments are shown; for (B) and (C) each biological replicate was conducted in technical triplicate and mean CFU values are shown.

**File S3. Full immunoblots and SDS PAGE analysis of the immunoblot samples for total protein content.** (**Pages 1-6**) Full immunoblots for Fig. 2A and S4B. On the left of each page, the relevant figure panel is shown and the lanes in question are marked with red outline. On the right of each page, the full immunoblot is displayed with the corresponding area also marked with red outline. (**Pages 7-9**) SDS PAGE analysis of the immunoblot samples for total protein content. In each page, the immunoblot in question is indicated (by “Fig. 2A” or “Fig. S4B”) and lanes are marked accordingly to identify the immunoblot lane that they correspond to (see white labels at the bottom of the gel).

**File S4. Analysis of *Stenotrophomonas spp.* for the presence of MCR proteins.** Hidden Markov Models built from validated sequences of MCR-like and EptA-like proteins were used for the identification of MCR-like analogues in a total of 106 complete genomes of the *Stenotrophomonas* genus downloaded from the NCBI repository. **(A)** Most genomes that were investigated (“*Stenotrophomonas maltophilia* genome” column”), encoded one or two MCR-like proteins (“Number of MCR analogues column”). **(B)** The 146 MCR-like sequences (“Protein ID column”) that were identified (only hits with evalues < 1e-10 were considered; “Evalue” column) belong to the same phylogenetic group as validated MCR-5 or MCR-8 proteins (“Phylogenetic group” column).

**File S5. Quality control information on 4,5-dibromo-2-(2-chlorobenzyl)pyridazin-3(2H)-one.** ^1^H-NMR and LCMS spectra of 4,5-dibromo-2-(2-chlorobenzyl)pyridazin-3(2H)-one (compound 36) demonstrating the correctness and purity of the synthesized compound by Bioduro-Sundia.

